# Aberrant splicing of *MBD1* reshapes the epigenome to drive convergent myeloerythroid defects in MDS

**DOI:** 10.1101/2025.10.17.682569

**Authors:** He Tian (Tony) Chen, Pratik Joshi, Severine Cathelin, Soheil Jahangiri, Joshua Xu, Emily Tsao, Yulin Mo, David Kealy, Adam Dowle, Saeer A Adeel, Zaldy Balde, Dylan Gowlett-Park, Katarina Czibere, Alexandra Misura, Olga Bigun, Renato Sasso, August Lin, Noor Kundu, Dianne Chadwick, Sila Usta, Tina Khazaee, Signy Chow, Hubert Tsui, Mark D Minden, Andrew N Holding, Katherine S Bridge, Gang Zheng, Kristin J Hope

**Author notes:** equal contributors (He Tian (Tony) Chen, Pratik Joshi). corresponding author **Corresponding author:** Kristin J. Hope, Princess Margaret Cancer Center, 101 College Street, Toronto, Ontario M5G 1L7, Canada. **Data sharing statement:** Raw and processed omics data have been deposited, and are available to view, on EGA (Study ID: EGAS50000000946, Dataset ID: EGAD50000001389). Pre-publication access for RIME proteomics data can be obtained and is available to view from MassIVE with the following link: ftp://, username = MSV000097058, password = o8R048Rk2Z41HVwp.

## Abstract

Myelodysplastic neoplasms (MDS) feature hematopoietic deficits driven in part by transcript splicing abnormalities. Thus far, such disease-driving transcripts have been identified in association with specific splicing factor mutations. However, it remains unclear whether there also exists a set of disease-wide conserved pathological transcripts, which drive MDS independently of mutational status. Here, we characterize an MDS-associated long isoform of *MBD1* (MBD1-L) as the first described member of this class of transcripts. Overexpression of MBD1-L in healthy human HSPCs recapitulates archetypal defects of MDS including deficits in erythroid differentiation and reconstitution capacity. These defects arise from an isoform-specific switching of MBD1’s binding behavior, refocusing its heterochromatin-promoting activity from methylated to unmethylated CpGs and enacting broad downregulation of CpG-rich promoters as well as secondary epigenetic effects mediated by its downstream target *BCOR*. Remarkably, we also find that directly reversing abnormal MBD1 splicing in primary human MDS using nanoparticle-encapsulated ASOs enhances erythroid differentiation.

**Key points:**

- Global mis-splicing of *MBD1* represents a novel gain-of-function epigenetic axis driving erythropoietic and proliferative defects in MDS.
- ASO based depletion of pathogenic MBD1 transcripts restores erythroid differentiation, advancing RNA-based therapies for MDS.

## Introduction

Myelodysplastic neoplasms (MDS) are age-associated clonal malignancies characterized by impaired hematopoiesis resulting from ineffective terminal cell maturation and dysfunctional commitment of hematopoietic stem cells (HSC)^1–7^. While ~30% of MDS cases evolve into highly aggressive secondary acute myeloid leukemia, MDS itself imposes significant disease burden^8, 9^ with non-AML mortality being the primary driver of survival outcomes across all risk groups^10–12^. Erythropoiesis is commonly impaired in MDS, leading to anemia and transfusion dependence. Thus, a major treatment goal for MDS is to mitigate transfusion dependence by targeting the MDS clone for differentiation using hypomethylating agents (HMAs), or by promoting hematopoiesis using erythropoietin analogs or Luspatercept. Such approaches however only yield an erythroid response in around half of all patients, with even fewer achieving transfusion independence^13–16^. Responses are also often transient, leaving few alternative options after failure of first-line agents. These clinical challenges underscore the need to attain a deeper understanding of the developmental and molecular underpinnings of MDS with a view towards identifying novel, effective treatment strategies.

Splicing dysregulation is a core feature of MDS, with 60% of cases exhibiting a recurrent splicing factor (SF) mutation in *SF3B1*, *SRSF2*, *U2AF1*, or *ZRSR2* ^17^. The resulting aberrant alternative splicing (AS) events contribute to hematopoietic dysfunction by altering the function or stability of key target transcripts^18–21^. All such AS events described so far however occur in a specific SF mutational subtype and are not found in other SF-mutant or SF-wildtype disease. However, splicing abnormalities can also arise by mutation-independent mechanisms^22^. Moreover, signatures of aberrant splicing have general prognostic power across MDS mutational subgroups, including SF wildtype^23^. These findings suggest that, in addition to mutation-specific splicing, the MDS state may also involve a parallel set of mutation-independent and potentially disease cross-cutting splicing events. If so, these events could highlight convergent disease-promoting mechanisms, representing promising global therapeutic targets. However, it remains unclear whether a shared class of alternatively spliced drivers truly exist in MDS, or whether SF-mutant and SF-wildtype MDS orchestrate their disease phenotype through distinct mechanisms.

Here, we identify and characterize splicing events shared between SF mutant and wildtype MDS and find a sizeable overlap of alternative splicing (AS) events between MDS mutational subtypes. We describe the AS of the epigenetic reader protein Methyl-binding-domain 1 (*MBD1*) as one such event. Instead of being instigated by SF mutation, we find that this abnormal MBD1 splicing is driven by reduced WTAP-m6A writer activity. Through broad rewiring of its chromatin binding, mis-spliced MBD1 distorts HSPC function to produce an isoform-specific impairment of erythropoiesis and response to hematopoietic demand, mirroring the sustained anemia and cytopenia seen in MDS. Remarkably, we find that reversing excess mis-splicing of MBD1 ameliorates MDS erythroid differentiation defects.

## Results

### A CXXC-3 containing isoform of *MBD1* is globally upregulated in MDS

To examine the potential for non-SF mediated contribution to MDS splicing dysregulation, we analyzed published RNA-seq of CD34^+^ HSPCs from MDS patients and healthy controls. Interestingly we found that splicing regulators are negatively enriched as a group in MDS regardless of SF mutant status, with *U2AF1* and *SRSF2* transcripts reduced in SF-wildtype MDS (**Supplementary figure 1A, B**)^24^. Using rMATs to identify AS events unique to MDS, we revealed >200 AS events displaying conservation across mutational subtypes including SF-wildtype (**Figure 1A**)^24, 25^. To enrich for AS events with functional involvement, we predicted alterations to transcript and protein sequence (**Figure 1B**)^26^. We accounted for impacts on translatable mRNA availability, such as altered CDS viability and nonsense-mediated decay (NMD), as well as frame-preserving changes affecting protein domains. Interestingly, we noted an enrichment for gain-of-function splicing changes among AS events, primarily through NMD rescue or by the inclusion of functional domains. (**Figure 1C**). Focusing on the latter as changes that could be amenable to therapeutic modulation, we identified inclusion of the frame-preserving alternative exon 11 (E11) in transcripts of *MBD1* as a key event of interest where preference for the E11-containing isoform (MBD1-L) over the E11-exclusion form (MBD1-S) is found in both SF-mutant and wildtype MDS with minimal change in total MBD1 abundance (**Figure 1D**).

**Figure 1:**
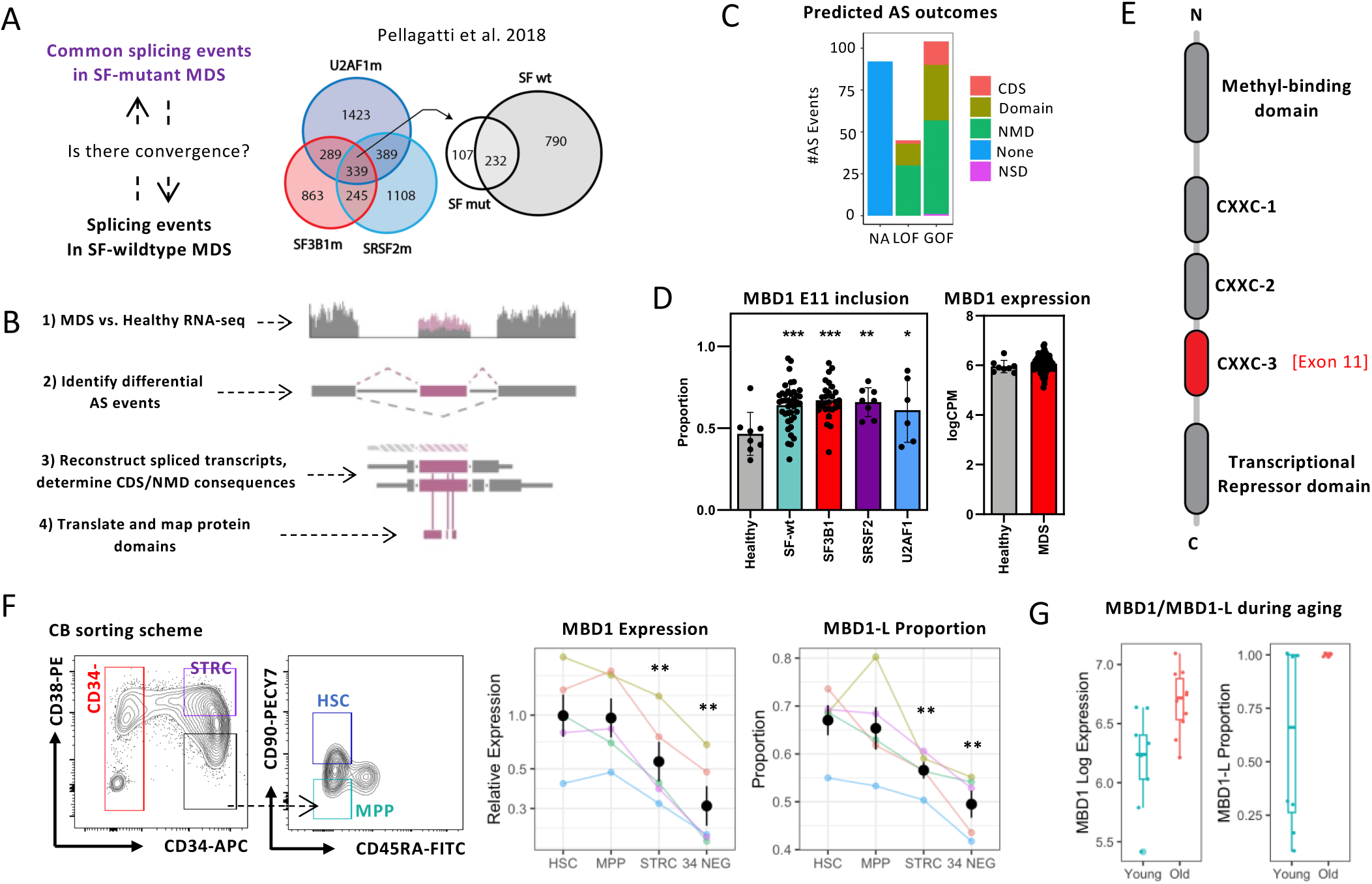
A CXXC-3 containing isoform of MBD1 is globally upregulated in MDS. **A** rMATs analysis using the Pellagati et. al. 2018 dataset reveals 232 splicing events enriched in CD34+ MDS HSPCs that are conserved between splice factor mutant and wildtype subgroups (82 MDS samples; 28 SF3B1 mutant, 8 SRSF2 mutant, 6 U2AF1 mutant and 40 splice factor wildtype). **B** Workflow for the identification of transcript alterations due to identified differential splicing events. **C** Predicted functional outcomes of MDS-wide AS events, by level of regulation. CDS: Establishment or disruption of a valid CDS. Domain: Alteration of an Interpro-annotated domain. NMD: Alteration in NMD sensitivity. NDS: Alteration in NSD sensitivity. **D** Proportion of MBD1-L in healthy controls or SF wildtype or mutant MDS subgroups, along with total MBD1 gene expression in healthy and MDS CD34+ HSPCs (healthy controls N=8). **E** Functional domains of MBD1. **F** qPCR of total MBD1 expression or MBD1-L proportion in CB CD34+38-90+45RA-HSCs, CD34+38-90-45RA-MPPs, CD34+38+ STRCs or terminal CD34-cells (N=5 samples). **G** MBD1-L proportion and total MBD1 expression in healthy young and aged CD34+38-HSPCs. (* p.val. ≤ 0.05, ** p.val. ≤ 0.01, *** p.val. ≤ 0.001)

MBD1 is a reader of methyl-CpG marks and establishes heterochromatin through recruitment of H3K9 methylase complexes. E11 contains MBD1’s CXXC-3 domain, which targets unmethylated CpGs, as opposed to the constitutive MBD domain that binds at methyl-CpGs (**Figure 1E**)^27–29^. CXXC-3 distinguishes MBD1 from all other MBD family members, which provide functional redundancy for repression of methylated, but not unmethylated, CpGs^30^. Although CXXC-3 is dispensable for MBD1’s repressive activity, its strong phylogenetic conservation likely portend its functional significance (**Supplementary figure 1C**)^31, 32^. Across the hematopoietic hierarchy, we found MBD1-L enriched in HSCs (CD34^+^CD38^−^CD90^+^CD45RA^−^) and MPPs (CD34^+^CD38^−^ CD90^−^CD45RA^−^), while progressively reduced in CD34^+^CD38^+^ progenitors and terminal CD34^−^ cells (**Figure 1F**). HSCs and MPPs also exhibited the highest expression of total MBD1 at RNA and protein levels (**Supplementary figure 1D**)^33–35^. Interestingly, MBD1-L increases further in these populations over the course of aging, where MDS incidence increases (**Figure 1G**)^36^.

### MBD1-L impairs the response to hematopoietic demand in an isoform-specific manner

We first determined whether MBD1-L imparts MDS-like defects to healthy human HSPCs by lentivirally overexpressing MBD1-L or -S in CB CD34^+^ HSPCs (**Figure 2A**, **Supplementary figure 2A**). We initially included a control for E11 sufficiency in the form of a second variant of MBD1-L with E13 excluded (MBD1-L*). *In vitro* competitive cultures showed a rapid depletion of MBD1-L-overexpressing cells, compared to a mild depletion for MBD1-S (**Figure 2B**). Importantly, MBD1-L and MBD1-L* produced identical phenotypes, indicating that CXXC-3 is specifically responsible for the observed defects. We additionally observed increased Annexin V, as well as impaired erythroid differentiation as seen by a decrease in CD71 with no change in myeloid differentiation (**Figure 2C,D**, **Supplementary figure 2B**). Colony forming assays showed decreased BFU-E and CFU-G colonies upon MBD1-L overexpression, with a magnitude more pronounced than for MBD1-S (**Figure 2E**). As differentiation defects were primarily in the erythroid lineage, and are prevalent in MDS, we performed erythroid differentiation cultures measuring erythroblast (EryII) and reticulocyte (EryIII) output. MBD1-L impaired terminal erythroid cell production by proportion and cell count, whereas MBD1-S increased the proportion of mature EryIII cells but with a mild decrease in cell count, implicating MBD1-L as a driver of dysfunctional erythropoiesis in MDS (**Fig. 2F**, **Supplementary figure 2C**).

**Figure 2:**
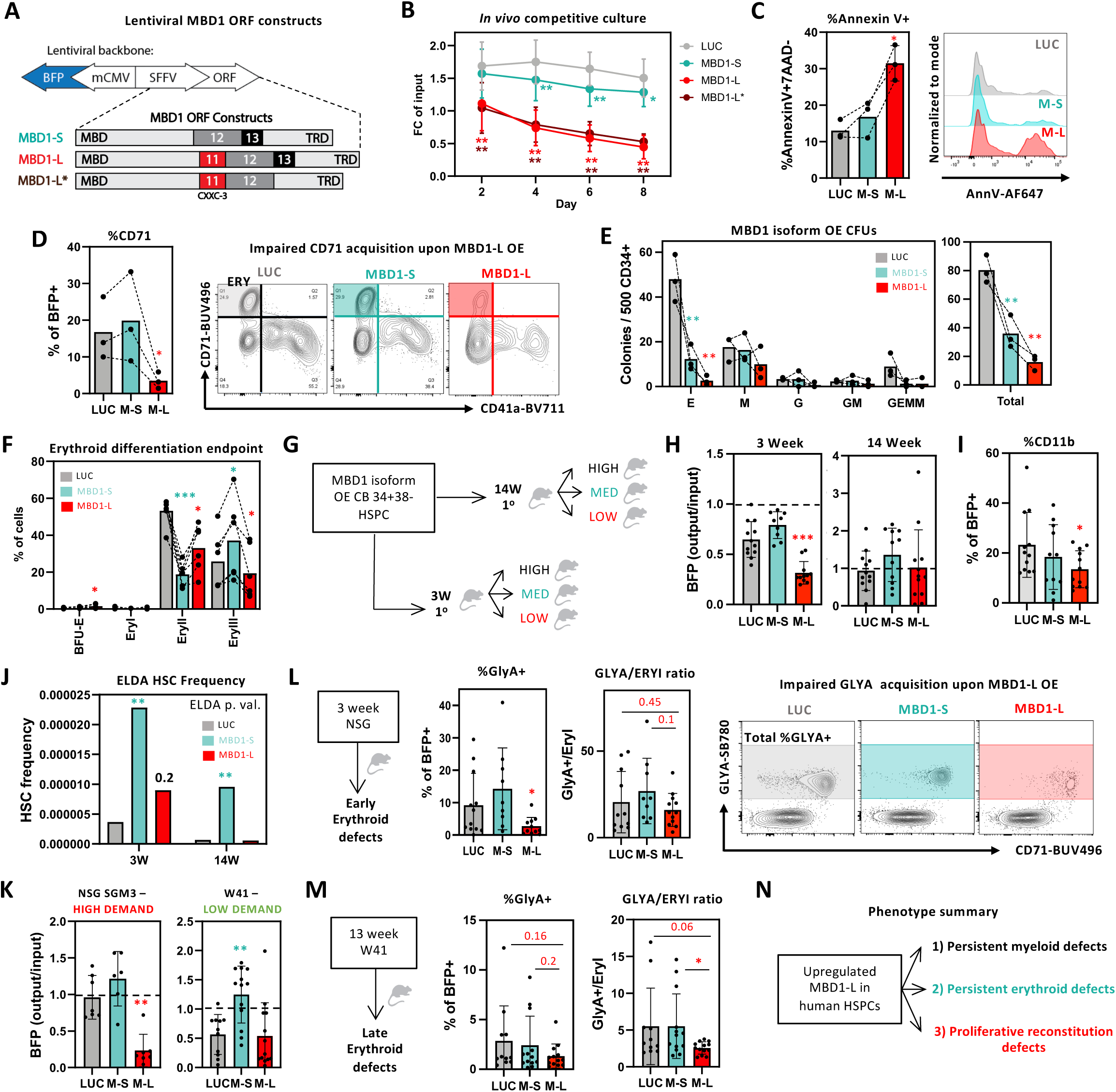
MBD1-L impairs the response to hematopoietic demand in an isoform-specific manner. **A** Schematic of MBD1 isoform lentiviral overexpression constructs. **B** *In vitro* competitiveness of cultured MBD1 isoform or control overexpressing CB Lin-CD34+ HSPCs. **C** *In vitro* Annexin V and 7AAD apoptosis analysis in transduced Lin-CD34+ HSPCs. **D** CD71 expression upon MBD1 isoform overexpression in Lin-CD34+ HSPCs. **E** Colony output of MBD1 isoform expressing HSPCs. **F** Proportion of erythroid lineage populations in MBD1 isoform overexpressing Lin-CD34+ HSPCs at the endpoint of a 2-step erythroid differentiation assay (N = 6 samples). **G** Schematic of 3-week and 14-week competitive transplants of isoform overexpressing Lin-CD34+38-HSPCs into NSG mice, followed by secondary transplants at limiting dilution. **H** Output/Input competition analysis within the injected femur of 3- and 14-week primary NSG xenografts. **I** CD11b proportion in primary 14-week NSG xenografts. **J** ELDA calculated HSC frequencies in pooled 3- and 14-week primary xenograft bone marrow. **K** Output/Input competition analysis within the injected femur of primary NSG-SGM3 and W41 transplants. **L** GlyA proportion and GlyA+/EryI ratio within the injected femur of 3-week primary NSG xenografts. **M** GlyA proportion and GlyA+/EryI ratio within the injected femur of primary W41 xenografts. **N** Summary of hematopoietic impairment upon MBD1-L overexpression in healthy HSPCs. N=3 samples for *in vitro* assays unless stated otherwise. All transplant data shown is from analysis of the injected femur unless stated otherwise. (* p.val. ≤ 0.05, ** p.val. ≤ 0.01, *** p.val. ≤ 0.001). All statistical tests are performed against Luciferase unless otherwise indicated.

We next assessed MBD1 isoforms *in vivo* using competitive xenotransplantation of CB Lin^−^ CD34^+^CD38^−^ HSPCs into NSG mice (**Figure 2G**). Tracking the proportion of transduced human cells over time revealed that MBD1-L hindered competitiveness during the proliferative initial phase of engraftment at 3 weeks but conferred no disadvantage in the steady state graft at 14 weeks (**Figure 2H**, **Supplementary figure 2E,F**). Nonetheless, we did observe reduced CD11b in late-stage MBD1-L grafts, highlighting persistent impairments in myeloid differentiation (**Figure 2I**). We then transplanted 3 and 14-week grafts into secondary recipients at limiting dilution to measure changes in HSC frequency and observed no difference between control and MBD1-L cells, suggesting that early MBD1-L competitive defects may primarily arise from reduced function, rather than loss, of HSCs (**Figure 2J**, **Supplementary figure 2G,H**). Interestingly, we observed a 7 and 11-fold increase in HSCs upon MBD1-S overexpression at 3 and 14 weeks respectively, supported by increased engraftment and CD34+ levels in secondary recipients of a non-limiting dose (**Supplementary figure 2I**).

To further explore MBD1-L’s reconstitution defects specifically at proliferative stages of reconstitution, we performed primary transplants into non-irradiated W41 mice, a low-demand environment with no irradiation-induced BM ablation, or irradiated NSG-SGM3 mice, where transgenic overexpression of human cytokines drives chronic proliferation. The graft-suppressing effect of MBD1-L persisted into the long-term in NSG-SGM3, but not W41 mice, suggesting that excess MBD1-L hinders the proliferative response during conditions of high hematopoietic requirement (**Figure 2K**).

Finally, we assessed impairments in erythroid differentiation *in vivo*. As erythroid output occurs in NSG mice at ~3 weeks, we assessed the erythroid graft present at this time and observed that MBD1-L but not MBD1-S compromised mature erythroid output (GlyA^+^) (**Figure 2L**). Given that W41 mice model long-term human erythropoiesis *in vivo*, we additionally assessed terminal erythroid output at a late 13 week timepoint. By measuring the ratio of terminal GlyA^+^ cells to GlyA^−^CD71^+^ progenitors we confirmed that MBD1-L-overexpressing HSPCs were compromised in their capacity to produce terminal erythroid cells in contrast to control and MBD1-S, with a trend towards total reduced GlyA (**Figure 2M**). Altogether, our results suggest that the gain of MBD1-L in MDS drives myeloid, erythroid and proliferative defects during times of hematopoietic demand, possibly contributing to the inability of MDS marrow to upregulate cell output in response to proliferative signals associated with cytopenia (**Figure 2N**).

### MBD1-L broadly represses promoter CpG islands

Since our results support a dominant function for CXXC-3, we hypothesized that CXXC-3 recruits MBD1-L’s heterochromatin-forming complex to new genetic loci inaccessible to MBD1-S^37^. We assessed the nuclear distribution of MBD1 isoforms via immunofluorescence microscopy on CB CD34^+^ HSPCs overexpressing MBD1-S or MBD1-L. Whereas MBD1-S localized near the nuclear periphery, rich in H3K9me3 heterochromatin, MBD1-L instead localized towards the nuclear interior (**Figure 3A**). This suggests that MBD1-S predominantly associates with constitutive heterochromatin whereas MBD1-L acts on transcriptionally active genic regions within the nuclear interior^38^. To map this in greater detail, we performed CUT&RUN on CB HSPCs overexpressing the MBD1 isoforms. Peak mapping onto published and unpublished enzymatic methyl-converted HSPC DNA methylation profiles revealed that MBD1-L prefers to bind CpG rich sites, whereas MBD1-S prefers methyl-CpGs, confirming that CXXC-3 actively shapes MBD1-L’s DNA binding footprint (**Figure 3B**) ^39^. In addition, the inclusion of CXXC-3 triggers a striking redistribution of MBD1-L to proximal promoters, likely driven by the presence of promoter-associated CpG islands (**Figure 3C**, **Supplementary figure 3A**). MBD1-L bound promoters were enriched in cell cycle-related genes, implicating cell proliferation as a likely control mechanism mediating MBD1-L’s hematopoietic defects (**Supplementary figure 3B**).

**Figure 3:**
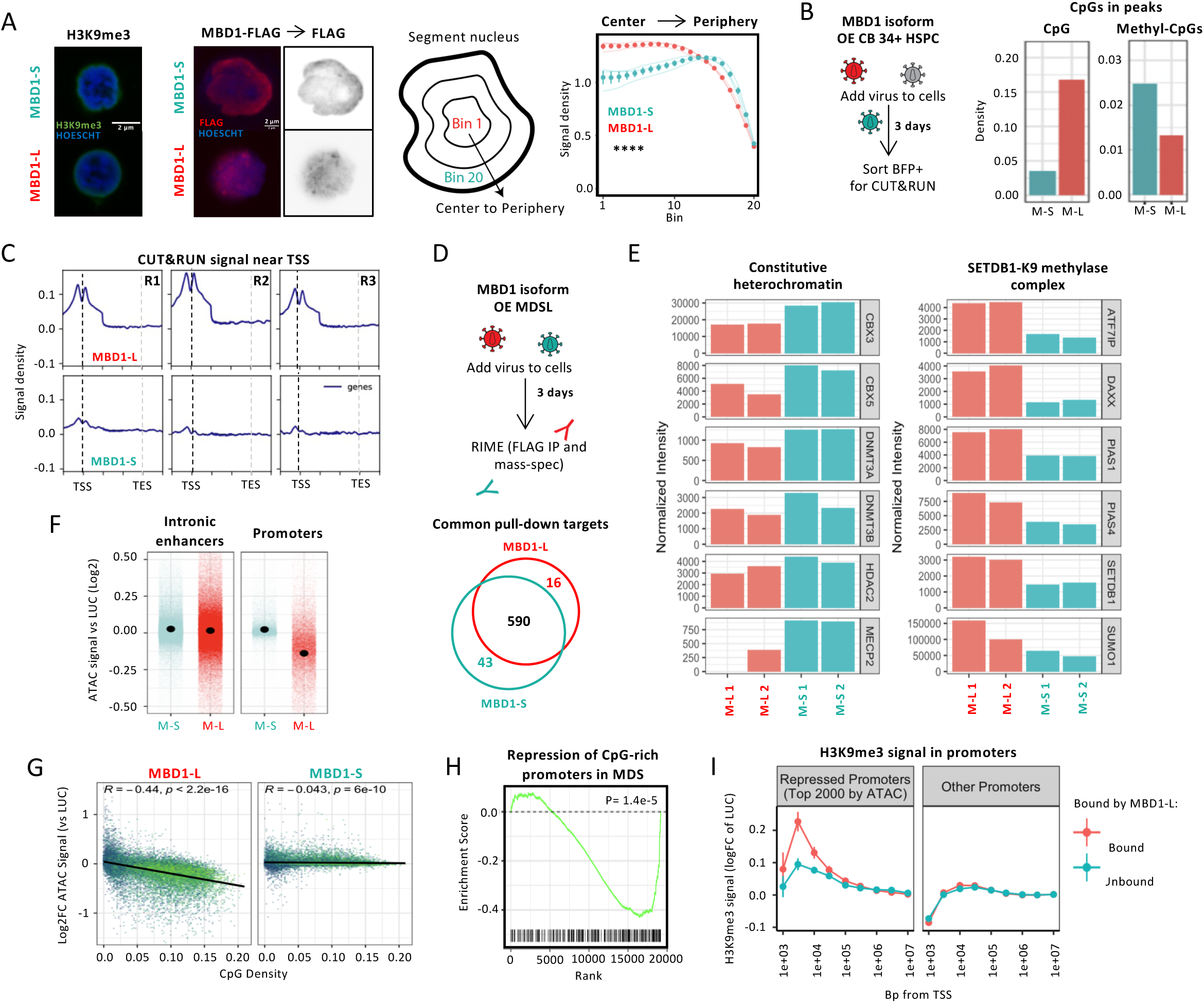
MBD1-L broadly represses promoter CpG islands. **A** H3K9me3 and MBD1 (via FLAG tag) staining in Lin-CD34+ HSPCs transduced with MBD1-L and MBD1-S lentivectors. FLAG staining was used to calculate the distribution of MBD1 isoforms within the nuclear area, measured 72 hours post-transduction (N = 30 analyzed cells per sample, with 3 samples used). **B** Estimated CpG and methylated CpG density within CUT&RUN peaks. **C** Peak distribution across gene regions in MBD1-L and MBD1-S CUT&RUN. **D** Schematic of RIME analysis of the MBD1-L and MBD1-S repressive complex in MDSL, with 590 targets overlapping between immunoprecipitated interacting proteins between the two isoforms. **E** Comparison of signal intensities of detected RIME-binding partners in MDSL. **F** Differential ATAC accessibility at intronic enhancers and promoters for MBD1 isoforms vs luciferase. **G** Correlation between ATAC peak repression by MBD1 isoforms and CpG density within the peak. **H** GSEA enrichment of the top 250 CpG-dense promoters in the Pellagati et. al. 2018 MDS Lin-CD34+ MDS HSPC RNA-seq. **I** H3K9me3 CUT&RUN signal proximal to the TSS at MBD1-L bound and repressed promoters compared to Luciferase. All multi-omics experiments were performed with 3 independent CB samples. (* p.val. ≤ 0.05, ** p.val. ≤ 0.01, *** p.val. ≤ 0.001)

We then asked whether the inclusion of CXXC-3 alters MBD1’s associated protein complex by performing Rapid Immunoprecipitation Mass Spectrometry of Endogenous Protein (RIME) profiling of chromatin-associated complexes^40^. The MBD1 interactome upon MBD1-L or MBD1-S overexpression in the MDSL cell line revealed that both isoforms associate with largely the same set of nuclear proteins (**Figure 3D**, **Supplementary figure 3C**). We did however observe quantitative differences in the level of association with bound proteins. MBD1-L exhibited an increased association with its canonical SETDB1 H3K9 methylase partner complex but exhibited reduced association with components of constitutive heterochromatin including the HP1 proteins CBX5 and CBX3, in agreement with the altered physical distribution of MBD1-L away from heterochromatic regions (**Figure 3E**)^41–45^.

To explore whether MBD1-L closes bound hypomethylated promoters we used ATAC-seq and observed that MBD1-L indeed produces global repression of ATAC signal at promoter regions, whereas MBD1-S does not. As expected, ATAC signal at intronic enhancers, which lack extensive CpG islands, showed no differences between MBD1-S and MBD1-L conditions (**Figure 3F**). Furthermore, the level of regional ATAC repression by MBD1-L correlated with its CpG density, indicating that the most CpG-rich promoters exhibit greater sensitivity to repression by MBD1-L (**Figure 3G**). Importantly, we identified that MDS HSPCs exhibit a negative enrichment of genes with CpG-rich promoters compared to healthy controls, suggesting that the observed pattern of MBD1-L repression is active in MDS (**Figure 3H**)^24^. Finally, we validated that the global closure of promoter regions by MBD1-L was associated with reduced transcription by measuring ethynyl uridine (EU) integration, and observed that MBD1-L inhibited the S-phase associated increase in transcription (**Supplementary figure 3D**).

Given that MBD1-L binds and represses promoter CpG islands, it stands to reason that these promoters would contain higher levels of repressive H3K9me3 marks. We found that overexpression of MBD1-L in HSPCs did not trigger an overall shift of H3K9me3 into the nuclear interior (**Supplementary figure 3E**) and reasoned that H3K9me3 deposition induced by MBD1-L may be more subtle. We thus profiled H3K9me3 and activating H3K4me3 histone marks at promoters using CUT&RUN on MBD1-S or MBD1-L overexpressing CD34^+^ HSPCs. Promoters containing MBD1-L binding peaks, representing direct loci of inhibition, indeed accumulated H3K9me3 marks within 1-10kb of the transcription start site upon MBD1-L overexpression (**Figure 3I**). Conversely, MBD1-L binding exhibited no correspondence with H3K4me3 status, confirming that MBD1-L’s closure of promoters depends on the installation of inhibitory H3K9me3 marks (**Supplementary figure 3F**).

### MBD1-L silences promoter networks to inhibit cycling and erythroid differentiation

To investigate downstream transcriptomic impacts of MBD1-L activity, we performed bulk RNA-sequencing. Sets of differentially expressed genes were highly distinct between MBD1 isoforms (**Figure 4A**). Additionally, repression of cell cycle genesets represented the major enrichment observed for MBD1-L, in accordance with the similar enrichment observed in its bound promoters profiled by CUT&RUN (**Figure 4B**). Indeed, ethynyl deoxyuridine (EdU)-labelled cell cycle assays revealed that MBD1-L impaired the transition of CB HSPCs from G1 to S phase (**Fig. 4C**). Importantly, this finding aligns with the observed downregulation of cell cycle pathways in MDS HSPCs (**Figure 4D**)^24^. Projection of the set of MBD1-L-repressed targets onto hematopoietic subpopulations revealed that MBD1-L downregulates genes associated with mature erythroid populations, consistent with our observed erythroid differentiation defects (**Supplementary figure 4A**) ^46^. However, we did not observe differential expression of key erythroid regulators, including *GATA1*, *GATA2*, *KLF1* and *TFRC*, suggesting that MBD1-L may regulate erythroid differentiation at the HSPC level via alternative means (**Supplementary figure 4B**).

**Figure 4:**
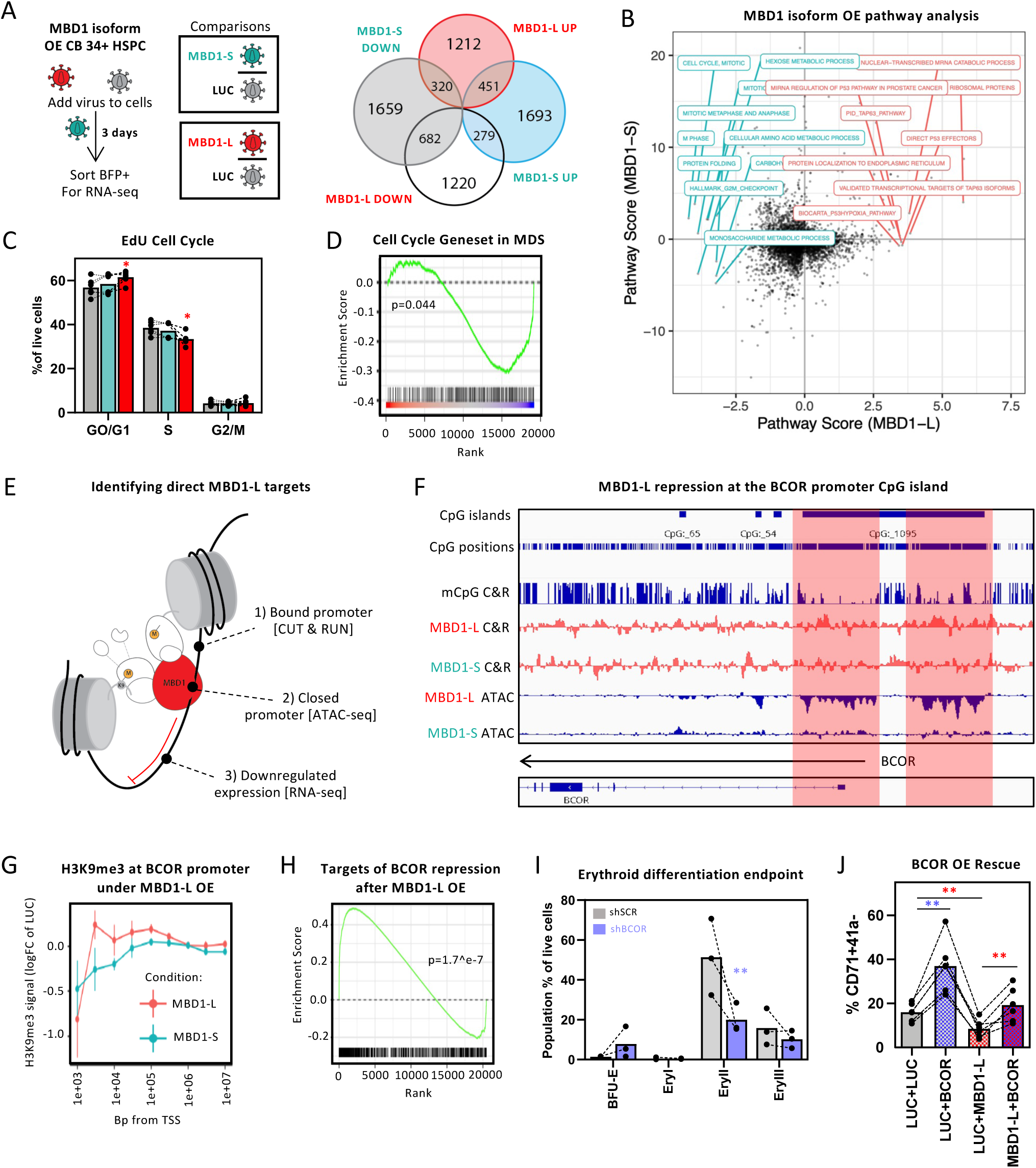
MBD1-L silences promoter networks to inhibit cycling and erythroid differentiation. **A** Overlap in differentially expressed genes between luciferase and either MBD1-S and MBD1-L transduced Lin-CD34+ HSPCs (p<0.05 vs Luciferase). **B** MBD1-L specific GSEA enriched terms. **C** Intracellular flow cytometry of MBD1 isoform overexpressing Lin-CD34+ HSPCs pulsed with EdU for 2 hours, measured 72 hours post-transduction (N=6 samples). **D** Cell cycle gene set enrichment in MDS CD34+ HSPCs (Pellagati et. al. 2018). **E** Schematic of pipeline used to select downstream direct targets of MBD1-L. Promoters bound by MBD1-L that were found to be less accessible, and their gene product was downregulated at the RNA level, were considered direct targets repressed by MBD1-L. **F** Tracks displaying MBD1-L and MBD1-S CUT&RUN and ATAC-seq signal in the BCOR promoter CpG island. CUT&RUN data is displayed as averaged peaks with IgG subtraction from the FLAG IP. ATAC-seq peaks are displayed as averaged logFE compared to Luciferase controls. CpG methylation status is also highlighted using an enzymatic methyl-converted CUT&RUN dataset. **G** H3K9me3 CUT&RUN signal proximal to the BCOR transcription start site in MBD1-L and MBD1-S overexpressing Lin-CD34+ HSPCs. **H** GSEA of BCOR-repressed targets in MBD1-L overexpression RNA-seq (bound targets obtained from Schaefer et al 2022). **I** Proportion of erythroid lineage populations in BCOR knockdown Lin-CD34+ HSPCs at the endpoint of a 2-step erythroid differentiation assay. **J** Rescue of the MBD1-L erythroid defect via BCOR co-expression. %CD71+41a-proportion in luciferase transduced Lin-CD34+ HSPCs that are either co-transduced with luciferase or a BCOR overexpression construct to establish baseline BCOR effects, or MBD1-L to establish the baseline erythroid defect, as well as Lin-CD34+ HSPCs transduced with MBD1-L and BCOR to measure restoration of erythroid differentiation (N=6 samples). All multi-omics experiments were performed with 3 independent CB samples. All *in vitro* assays are performed with 3 samples unless otherwise stated. (* p.val. ≤ 0.05, ** p.val. ≤ 0.01, *** p.val. ≤ 0.001).

To identify MBD1-L targets representing novel candidate effectors of our erythroid phenotype we integrated our omics datasets (**Figure 4E**), filtering for genes exhibiting MBD1-L-unique binding, ATAC-seq repression, and RNA-seq downregulation at a threshold of p<0.05. Among this set of direct targets was BCL6 corepressor (*BCOR)* (**Supplementary figure 4C**). BCOR participates in epigenetic gene silencing as part of the BCL6 repressive complex and Polycomb Repressive Complex 1.1, and its function is impaired in MDS by both recurrent mutations and expression downregulation (**Supplementary figure 4D**)^47,48,49^. The BCOR major isoform promoter is particularly notable as it is encompassed by a 15.8kb CpG island, the longest among protein-coding genes, making it a particularly sensitive site of regulation by MBD1-L (**Supplementary figure 4E**). This island features a methylated center flanked by hypomethylated regions corresponding to ATAC peaks. MBD1-L binds these flanking regions and suppresses the ATAC signal at these positions accompanied by an increase in H3K9me3 (**Figure 4F, G**). Furthermore, MBD1-L derepressed BCOR-controlled genes (**Figure 4H**). Several of the most upregulated genes upon MBD1-L overexpression are BCOR targets, including *GDF15*, a TGF-β ligand associated with ineffective erythropoiesis in MDS (**Supplementary figure 4F**)^50–52^. These results support a second level of indirect epigenetic regulation by MBD1-L, whereby de-repression of BCOR targets drives upregulation of a subset of genes in parallel with the broad promoter repression directly exerted by MBD1-L’s repressor scaffolding activity.

We next aimed to determine whether BCOR loss orchestrates the erythroid defects associated with excess MBD1-L. Two-step erythroid differentiation culture revealed lowered erythroblast (EryII) output in response to BCOR knockdown in CD34+ HSPCs (**Figure 4I**, **Supplementary figure 4G**). In contrast, rescue of BCOR levels in MBD1-L-overexpressing HSPCs by simultaneous transduction of a BCOR overexpression lentivirus completely rescued the MBD1-L-induced erythroid defect but yielded no improvement of the proliferative defect (**Figure 4J**, **Supplementary figure 4H,I**). This suggests that the proliferative and erythropoietic aspects of MBD1-L’s effects are independent phenotypes arising from parallel but distinct mechanisms with the loss of proliferation resulting from MBD1-L’s direct repression of cell-cycle related promoters, and the erythroid deficit instead attributable to secondary epigenetic repercussions stemming from BCOR insufficiency.

### The m6A writer WTAP regulates the splicing of MBD1-L

Since MBD1-L splicing is not associated with specific SF mutations, we hypothesized that RNA regulators globally altered across MDS might be responsible for pathological MBD1 splicing. We therefore scored for differentially expressed RNA binding proteins (RBPs) from MDS CD34^+^ transcriptomes whose expression correlated or anti-correlated with the inclusion of E11^24^. Intriguingly, of the group of RBPs deficient in MDS, *WTAP*, a component of the N6-methyladenosine (m6A) writer complex, exhibited the strongest negative correlation with E11 inclusion (**Figure 5A**, **Supplementary figure 5A**). We also found that WTAP was downregulated in CD34^+^ MDS HSPCs, consistent with increased MBD1-L (**Figure 5B**). Supporting WTAP’s role in regulating MBD1-L splicing, we found that its knockdown increased the MBD1-L proportion in MDSL cells. Likewise, knocking down METTL3 and METTL14, the two other core members of the m6A writer complex, also increased MBD1-L splicing, suggesting that although MBD1-L splicing in MDS is primarily associated with WTAP depletion, any loss in m6A writer function also biases towards the MBD1-L isoform (**Figure 5C**, **Supplementary figure 5B**). Furthermore, inducing a WTAP-depleted state in normal CB HSPCs via knockdown recapitulated the proliferative and erythropoietic defects induced by MBD1-L overexpression (**Figure 5D,E**). These results overall support reduced m6A marks, facilitated by reduced WTAP expression, as a key mechanism enforcing inclusion of E11. Interestingly, we also identified a large-scale correlation between the splicing signature induced by knockout of m6A writers *WTAP* and *METTL14* and the MDS splicing signature, suggesting that m6A modifications may be implicated more broadly in the perturbed splicing state of MDS (**Supplementary figure 5C**)^19, 53^.

**Figure 5:**
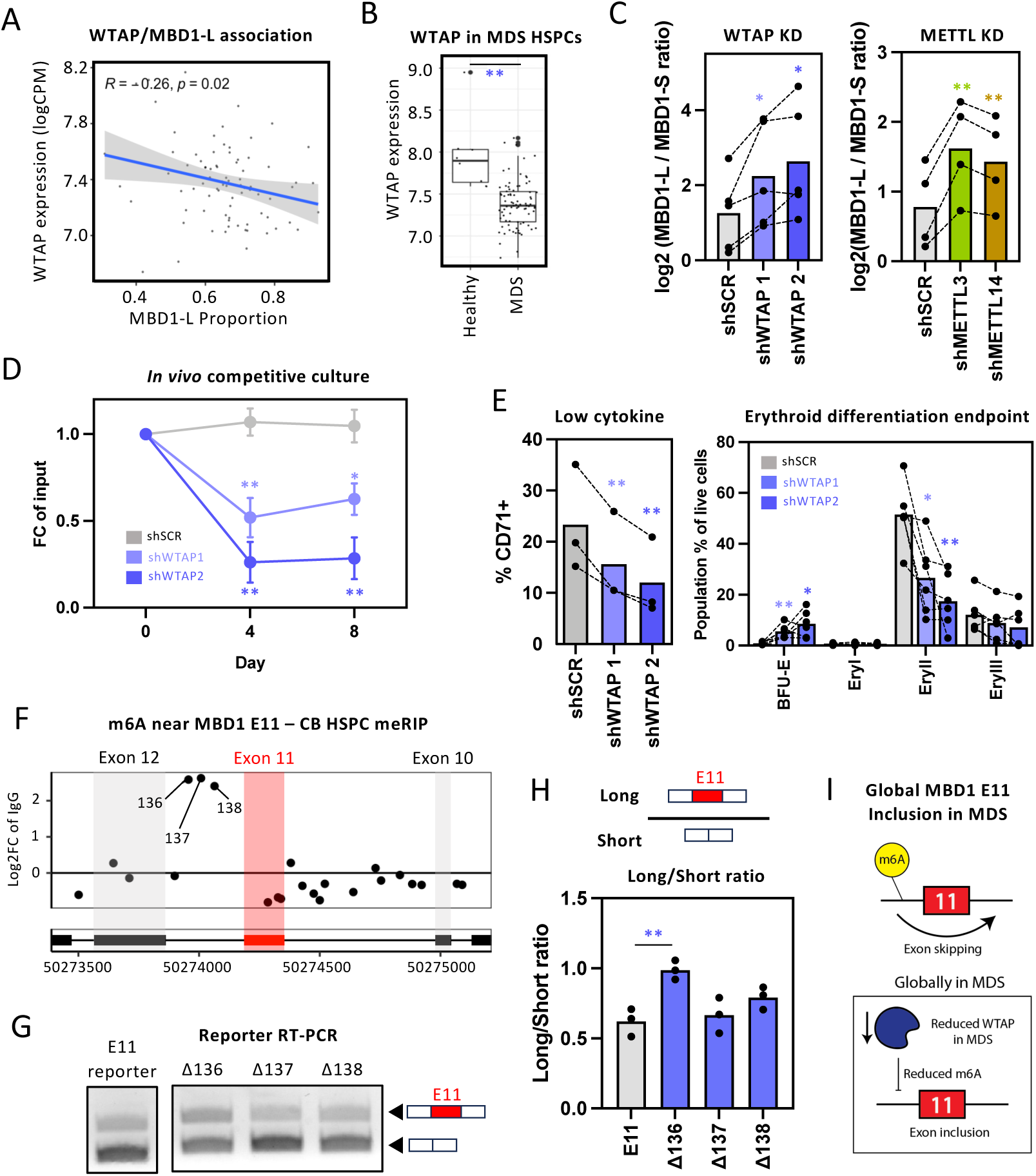
The m6A writer WTAP regulates the splicing of MBD1-L. **A** Correlation between MBD1-L inclusion and WTAP expression in MDS CD34+ HSPCs (Pellagatti et. al. 2018). **B** Expression of WTAP in healthy versus MDS CD34+ HSPCs (Pellagatti et. al. 2018). **C** qPCR quantification of MBD1-L/S ratio 6 days post-transduction with shRNAs against WTAP (N=5 biological replicates), or METTL3 and METTL14 (N=4 biological replicates) in MDSL. **D** *In vitro* competitiveness of cultured Lin-CD34+ HSPCs transduced with shRNAs against WTAP. **E** CD71 proportion in regular cytokine culture upon WTAP knockdown (N=3 samples) and proportion of erythroid lineage populations in WTAP knockdown Lin-CD34+ HSPCs at the endpoint of a 2-step erythroid differentiation assay (N=6 samples). **F** MeRIP peak enrichment at DRACH motifs along the MBD1 transcript in CB CD34+ HSPCs (Li et. al., 2018). **G** Representative RT-PCR of mini-spicing reporter derived *MBD1* transcripts in MDSL. **H** E11 mini-splicing reporter quantification of the ratio between the E11 inclusion and exclusion forms. **I** Proposed mechanism of global increase in MBD1 E11 inclusion in MDS. All functional assays are performed with 3 samples unless otherwise stated. (* p.val. ≤ 0.05, ** p.val. ≤ 0.01, *** p.val. ≤ 0.001).

The correspondence between loss of m6A activity and increased E11 inclusion suggests that m6A modification promotes E11 skipping. To identify the specific sites responsible for this regulation, we referenced published CB CD34+ HSPC meRIP data and calculated meRIP peak enrichment at DRACH motifs. This analysis identified 3 candidate m6A-marked motifs within the E11 downstream intron (sites 136, 137 and 138) (**Figure 5F**) ^54, 55^. To test whether m6A disruption within these sites could increase E11 inclusion, we created splicing minigene reporters spanning E10 to E12, with the target m6A sites inactivated by A>T point mutation. We transduced MDS-L cells with these reporters and performed RT-PCR using reporter-specific primers to assess E11 inclusion levels (**Figure 5G**). Of the three sites tested, the Δ136 mutation significantly increased the proportion of spliced reporter transcripts containing E11, suggesting that loss of m6A marks at this location due to downregulation of WTAP in MDS drives the pattern of preferential MBD1-L splicing across MDS (**Figure 5H,I**).

### Depletion of *MBD1* restores differentiation in MDS cells

Given that excess MBD1-L imposes significant hematopoietic impairment, we asked whether reducing MBD1-L can rescue the defective hematopoietic output and erythroid differentiation seen in MDS cells. Because MBD1 disruption in MDS affects solely splicing and not total MBD1 abundance, we elected to directly modulate E11 splicing to simulate restoration of a healthy splicing state while maintaining normal MBD1 levels by using an antisense RNA oligonucleotide (ASO) which sterically inhibits SF recognition of the 3’ splice acceptor site of E11 and prevents E11 inclusion (**Figure 6A**)^56^. To optimize uptake and delivery, we packaged E11-targeting ASOs into lipid nanoparticles (LNPs) using the current benchmark formulation for LNP-based small RNA delivery^57^. Testing a variety of nanoparticle-payload proportions calibrated by amine/phosphate (N/P) ratio to mediate delivery of RNA cargo into CD34^+^ HSPCs, we found the N/P ratio of 12:1 produced efficacious RNA delivery, through detection of intracellular FAM-conjugated control RNAs, as well as MBD1 splicing modulation in MDS cells upon E11 targeting ASO treatment (**Figure 6A**, **Supplementary figure 6A**).

**Figure 6:**
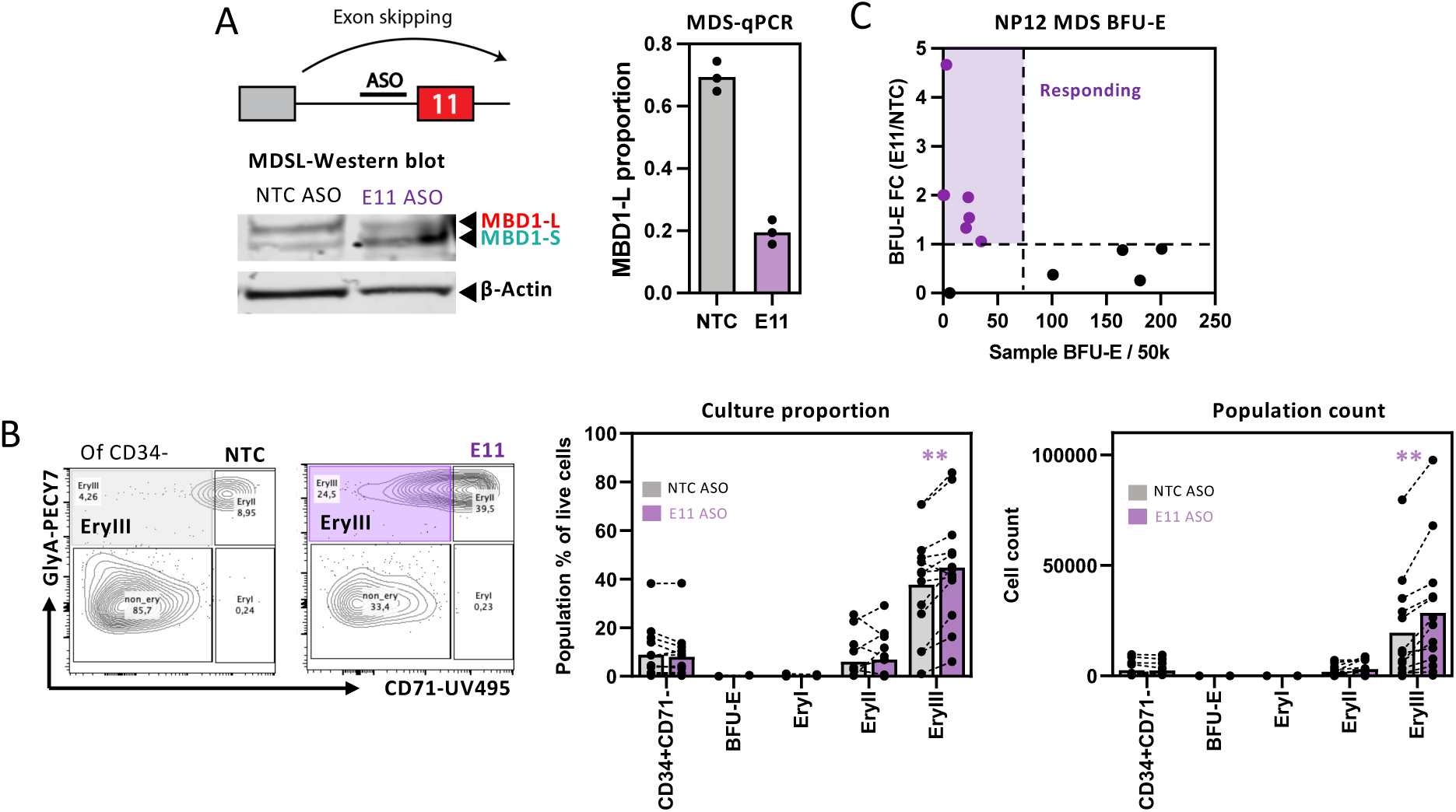
Depletion of MBD1 restores differentiation in MDS cells. **A** Schematic of E11 targeting ASOs impeding E11 inclusion. MBD1-L proportion measured via qPCR after a pulse of control (NTC) or E11 ASO in MDS CD34+ HSPCs using ASO packaged in LNPs with a N/P ratio of 12:1 (N=3 technical replicates). Additional ASO validation was performed using a western blot of MBD1-L and MBD1-S in MDSL cells treated with control and E11 ASOs. **B** Restoration of terminal EryIII proportion and absolute count in primary MDS CD34+ HSPCs treated with E11 or control ASOs at the day 18 endpoint of a 2-step erythroid differentiation assay with MDS HSPCs cultured on a primary human bone marrow mesenchymal stromal cell layer (N=13 samples). **C** BFU-E colonies of primary MDS cells pulsed with E11 or control ASO. Fold change in BFU-E colonies compared to the NTC control are plotted against total BFU-Es in NTC samples. Samples with a BFU-E fold change greater than one are highlighted as responding samples (N=11 samples). All functional assays are performed with 3 samples unless otherwise stated. (* p.val. ≤ 0.05, ** p.val. ≤ 0.01, *** p.val. ≤ 0.001).

To assess the therapeutic potential of the validated E11 ASO, we isolated CD34^+^ MDS HSPCs from 13 treatment naïve patient samples and cultured cells on human BM mesenchymal stromal cells (MSCs), treating with ASO at 0, 6, and 12 days of 2-step erythroid culture. Remarkably, we found that the E11 ASO increased the proportion and number of late erythroid cells at the culture endpoint across the majority of tested samples compared to the non-targeting (NTC) ASO (**Figure 6B**, **Supplementary figure 6B,C**). We likewise assessed erythroid progenitor function on MDS cells from 12 patients after a 24-hour ASO treatment. This brief ASO treatment yielded an increased erythroid colony output specifically for samples that displayed low basal erythroid output, suggesting MBD1-L depletion may be most beneficial for those cases exhibiting severe erythropoietic impairment (**Figure 6C**). Separately, we observed a similar increased erythroid colony output in MDS samples with impaired basal erythropoiesis using a less efficacious ASO formulation tested in an additional 10 primary samples, further supporting the E11 ASO therapeutic potential (**Supplementary figure 6D**).

## Discussion

We pursued the novel approach of searching for common underlying driver AS events that contribute to the MDS disease state, and uncovered MBD1-L as a key pathogenic isoform. MBD1-L’s repressive action at promoters inhibits cell cycle processes to reduce hematopoietic output, while the specific repression of the BCOR promoter impairs erythroid differentiation. As cycling rate regulates erythroid commitment and maturation, impaired cell cycling may additionally contribute to the MBD1-L-induced erythroid deficit^58^. Dysregulated cell cycling and increased apoptosis are hallmark features of ineffective erythropoiesis in MDS, and the dampening of responsiveness to proliferative signaling by MBD1-L-overproduction may underlie the marrow’s deficient response to anemic stress^59–61^.

Interestingly, similar abnormalities in erythroid and cell cycle processes are observed in murine models of *Tet2* haploinsufficiency. Since *TET2* depletion represses promoters by impairing the erasure of DNA methylation, these phenotypes may represent conserved sequelae of aberrant CpG island repression in MDS^62^. In addition, we observed an isoform switch from MBD1-S to -L in HSPCs with aging, where our findings suggest that MBD1-S is responsible for reinforcement of constitutive heterochromatin, and this serves a reconstitution-enhancing function. Increased MBD1-L with aging may degrade the capacity to maintain organized heterochromatin, driving functional decline and possibly increasing the risk of leukemogenesis. Indeed, disruptions to the abundance and spatial distribution of H3K9me3 are recurrently observed in HSPCs upon aging and can contribute to aging phenotypes^63–65^. To this point, we found that MBD1-L lacks spatial association with heterochromatin in the nuclear periphery and instead migrates to central genic regions where H3K9me3 marks spread to active promoters with low CpG methylation. Our work thus showcases MBD1-L’s preferential production with age and dominant capacity to alter epigenetic landscapes and function in ways that mirror aged and/or pre-leukemic HSPCs.

MBD1-L represents to our knowledge the first described instance of m6A-mediated pathogenic splicing in MDS. Loss of the WTAP complex has been shown to promote the inclusion of GC-rich exons flanked by short introns, all characteristics of MBD1 E11.^66^ Our work additionally revealed a remarkable correspondence between the MDS splicing signature and global splicing alterations upon *WTAP* and *METTL14* knockout. These findings point to m6A modifications as a potentially unifying mechanism for AS dysregulation in MDS, wherein the effects of individual SF mutations are superimposed onto a background of splicing disruption. Further characterization of the extent of this regulatory relationship could have far reaching impacts on our understanding of splicing disruption in MDS and its treatment. This extends to other hematological malignancies, including AML, or solid cancers where SF mutations have been observed, yet have a low mutation rate that restricts availability of gene-level targets^67^.

In a first-time application of ASOs in MDS, we show that re-equilibration of the MBD1-L/MBD1-S ratio in patient samples results in a therapeutic increase in erythroid output. A limited 24-hour exposure of MDS HSPCs to the ASO was also sufficient to ameliorate impaired erythropoiesis at the level of progenitors in patients with particularly impaired erythropoiesis, demonstrating the powerful effects of MBD1-L targeting ASOs and sets the stage for further exploring this approach in MDS therapeutics independently or in synergy with interventions targeting terminal red cell maturation, such as luspatercept^4, 15, 61^. As MBD1-L acts in a methylation-independent manner, we expect that any deleterious effects resulting from its epigenetic influence would be intractable by hypomethylating agents currently used as first-line MDS therapies. A corollary of this is that increased MBD1-L splicing may be a contributory mechanism to HMA-refractory anemia in MDS and could potentially be an indicator for poor HMA response, whose predisposing factors are not currently well understood.

Our work highlights that in contrast to the consensus view, an extensive landscape of aberrant AS events is shared across MDS containing a subset of gain-of-function events which represent a novel, untapped pool of potential therapeutic targets. We demonstrate the therapeutic utility of this class characterizing an AS event in MBD1, which through broad epigenetic remodeling impairs HSPC-driven erythropoiesis and response to hematopoietic demand. That ASO mediated correction of MBD1 splicing can independently drive increased erythroid output in MDS patient samples highlights this as a novel MDS therapeutic strategy well positioned for further development.

## Acknowledgements

We would like to thank the Princess Margaret Flow Cytometry Core, Princess Margaret Genomics Center, SickKids Center for Applied Genomics and the University Health Network Animal Research Center. This work was supported by a Canadian Institutes of Health Research (CIHR) Doctoral Scholarship (H.T.T Chen), a Natural Sciences and Engineering Research Council of Canada (NSERC) Doctoral Scholarship (E. Tsao), a CIHR Master’s scholarship and an Ontario Graduate Scholarship (P. Joshi), a Princess Margaret Cancer Foundation Catalyst Grant (to K.J. Hope), a Cancer Research Society grant (to K.J. Hope) and an Ontario Institute for Cancer Research (OICR) Investigator Award (K.J. Hope).

## Author Contributions

H.T.T.C and P.J designed and performed experiments, analyzed data, interpreted results and wrote the manuscript. S.C, J.X, E.T, S.A.A and Z.B designed and performed experiments, as well as analyzed data and interpreted results. D.K and A.D, under the supervision of K.S.B and A.N.H, performed and analyzed the MBD1 isoform RIME. Y.M, under the supervision of G.Z, generated and performed quality control on the MBD1 ASO LNPs. D.G.P, K.C, A.M, O.B, R.S, A.L, N.K, D.C, S.U, T.K and S.C, under the supervision of H.T, sourced and banked primary MDS samples. M.D.M additionally provided primary MDS samples. K.J.H conceived of and supervised the project, designed experiments, interpreted results and wrote the manuscript. All authors reviewed the manuscript. The authors declare no conflicts of interest.

## Methods

### Primary cell isolation and culture

Informed consent was obtained from all primary tissue donors, and all work was conducted in accordance to the Research Ethics Boards at University Health Network (UHN) Research Ethics Board (CAPCR # 20-6026) in accordance with Canadian Tri-Council Policy Statement on the Ethical Conduct for Research Involving Humans (TCPS). Umbilical cord blood was obtained from healthy donors with uncomplicated deliveries. Cord blood lineage depleted cells were isolated using Ficoll separation followed by lineage depletion using EasySep Human Progenitor Cell Enrichment Kit II (StemCell Technologies, Cat. #19356) as previously described^68^. Cord blood CD34+ cells were purified by flow sorting and cultured in StemSpan SFEM II (StemCell Technologies, Cat. #09655), supplemented with 100ng/mL SCF, 100ng/mL FLT3-L, 20ng/mL IL-6, and 20ng/mL TPO. Lentiviral transductions of HSPCs were performed after >12h of recovery. Primary MDS cells were obtained from diagnostic bone marrow aspirates, provided by the Sunnybrook Hematology Biobank, collected prior to initiation of disease-modifying treatment. Bone marrow aspirates were collected in sodium heparin or EDTA collection tubes and MNCs are isolated using gradient centrifugation on Ficoll-Paque. MDS CD34+ cells were isolated as previously described^69^, using the MACS CD34+ enrichment kit (Miltenyi, Cat. # 130-046-702) and cultured in StemSpan SFEM II supplemented with 100ng/mL SCF, 100ng/mL FLT3-L, 20ng/mL IL-6, 20ng/mL TPO, and 10ng/mL IL-3. Stromal cell supports were prepared using human BM mesenchymal stromal cells (MSC), obtained from femoral marrow aspirates from hematologically healthy individuals undergoing orthopedic surgery, grown in ɑ-MEM supplemented with 1x GlutaMAX, 2U/mL sodium heparin and 10% human platelet lysate (StemCell Technologies, Cat. #06960). The stromal media was replaced with MDS media prior to seeding. For erythroid differentiation culture, CB CD34+ HSPCs were plated in SFEM II with Stemspan Erythroid Expansion Supplement (StemCell Technologies, Cat. #02692). After a 6-day culture, cells were then switched to SFEM II with 3U/mL EPO, 2U/mL sodium heparin and 3% human platelet lysate to induce maturation for up to day 16. For MDS CD34+ HSPCs, the media change was performed at 6 days. Colony forming unit (CFU) assays were performed in ColonyGEL Human Complete Medium (ReachBio, Cat. #1102). 750 cord blood CD34+ or 25,000 MDS MNCs were plated in 1.5mL media and cultured for 12-16 days prior to colony scoring.

### Lentiviral vectors

C-terminal FLAG or V5 tagged MBD1 isoform open-reading frames, as well as the native BCOR open reading frame, were lentivirally overexpressed under the SFFV promoter using the pSMALB backbone, a gift from Drs. John Dick & Peter van Galen (Addgene ##161786, RRID: Addgene_161786). shRNAs were lentivirally expressed using pLKO.1-GFP, cloned in house from pLKO.1-puromycin by replacing the puromycin resistance cassette with eGFP. pLKO.1-puromycin was a gift from Dr. Bob Weinberg (Addgene ##8453). shRNAs were selected from the Broad TRC shRNA Library and validated by qPCR. The control shScramble random-mer sequence was derived from PLKO.1-shScramble (Sabatini Lab, Addgene ##1864). To clone E11 mini-splicing reporters, wildtype or A > T mutated Δ136 gBlocks (IDT) were ordered and cloned into pSMALb. Δ137 and Δ138 A > T mutations were made using the wildtype gBlock template and site-directed mutagenesis.

### Quantification of MBD1 isoforms

RNA was isolated using the PicoPure RNA isolation kit (ThermoFisher, Cat. #KIT0204), reverse-transcribed using qScript cDNA SuperMix (Quanta Biosciences, Cat. #CA101414-106). The ratio of MBD1-L and MBD1-S was measured by qPCR using Taqman Fast Advanced qPCR Mastermix (ThermoFisher, Cat. #4444964) with primer sets spanning the exon 11 upstream splice junction and the exon 11-skipping junction. The reference gene was *ACTB*. All biological replicates were run in technical duplicates or triplicates. For RT-PCR, cDNA was amplified using OneTaq Quick-Load 2X Master Mix (NEB, Cat. #M0486S) using primers complementary to the upstream and downstream flanking exons.

### Cell culture and lentivirus production

LentiX-HEK293T cells (Takara Bio) were cultured in DMEM with 10% FBS and 1mM sodium pyruvate. Lentiviral transfer plasmid was transiently transfected using Lipofectamine 2000 transfection reagent (ThermoFisher, Cat. #11668500) alongside pMD2.G and psPAX2 packaging plasmids (Trono Lab, Addgene ##12259; ##12260) to create VSV-G pseudotyped lentiviral particles. Lentivirus was harvested 48-72h post-transfection and concentrated via ultracentrifugation (25,000g, 2h, 4°C), then resuspended in StemSpan SFEM II (Stemcell Technologies). The concentration of transduction units in lentiviral preparations was quantified through titration on HeLa cells (RRID: CVCL_0030). HeLa cells (RRID: CVCL_0030) were cultured in DMEM with 10% FBS. K562 cells (RRID: CVCL_0004) were cultured in RPMI with 10% FBS. MDS-L cells were cultured in RPMI with 10% FBS and 20ng/mL IL-3. MDS-L cells were a gift from Dr. Kaoru Tohyama.

### Antisense Oligonucleotides

A 2’-O-ethoxy-ethyl (2’MOE) modified splice-modulating antisense oligonucleotide (ASO) was designed with complementarity to the 3’ splice acceptor site of MBD1’s CXXC-3 exon. ASOs were encapsulated in lipid nanoparticles (LNPs) using an N/P ratio of 6:1 or 12:1. Cells were treated with the equivalent of 0.15uM ASO. LNPs were prepared using the microfluidic rapid mixing method as previously reported^70^. Lipids were mixed in ethanol at a molar ratio of 50/10/38.5/1.5 (DLin-MC3-DMA/DSPC/Cholesterol/DMG-PEG2000). ASOs were dissolved in 25 mM sodium acetate buffer (pH = 4.0). The amount of ASO was determined based on a 6:1 molar ratio between the total ionizable nitrogen groups in DLin-MC3-DMA and the overall negative charges from ASO. The two phases were mixed through herringbone microfluidic chips (microfluidic ChipShop, Germany) at a volumetric flow rate ratio of 3:1 (aqueous to ethanol) and total flow rate of 10 mL/min. The mixed solution was dialyzed against PBS 7.4 overnight. Afterwards, LNPs were concentrated using an Amicon 100 kDa MWCO centrifugal unit (Amicon®, Sigma-Aldrich, Cat. # UFC510008) and passed through 0.22 um filter before use. ASO concentration and encapsulation efficiency were measured by Quant-it™ RiboGreen RNA Assay (Thermofisher, Cat. #R11490).

### Flow Cytometry Assays

For immunophenotyping, cells were stained in flow buffer (PBS, 2% FBS, 2mM EDTA) using human CD45-FITC (HI30, BD Biosciences Cat. #555482), mouse CD45-APC (30-F11, BD Biosciences Cat. #559864), CD34-APC (BD Biosciences Cat. #555824), CD33-BUV395 (P67.6, BD Biosciences Cat. #745709), CD19-PECY7 (HIB19, BD Biosciences Cat. #560728), CD11b-PE (BD Biosciences Cat. #557321), CD71-BUV496 (M-A712, BD Biosciences Cat. #749247), CD235a-SB780 (GA-R2, eBioscience Cat. #78-9987-42), and CD235a-PECY7 (GA-R2, BD Biosciences Cat. #563666). Xenograft cells were blocked with mouse FC block (BD Biosciences, Cat. #553142) and human IgG prior to staining. For apoptosis analysis, cells were stained with Annexin V-AF647 (Thermofisher, Cat. #A23204) and 7-AAD (BD Biosciences, Cat. #559925) in Annexin Binding Buffer (BD Biosciences, Cat. #556454); surface markers were stained simultaneously. For Click-iT chemistry based assays, EdU, EU, and OPP labelled cells were first fixed with 4% paraformaldehyde (PFA) for 25 minutes, followed by permeabilization with 0.5% Triton-X and labelling using the Click-iT flow cytometry assay kit (ThermoFisher, Cat. #C10269). The labelling conditions were: EdU-10nM for 2h, EU-1uM for 1h, and OPP-20uM for 1h. Analysis was done using FlowJo (RRID: SCR_008520).

### Immunofluorescence Microscopy

Cells were spun onto Poly-L-lysine coated Ibidi u-Slide 18-well glass bottom chambered slides (Ibidi, Cat. #81817) at 200xg for 10 minutes, fixed with 4% PFA in PBS for 20 minutes and permeabilized with 0.1% Triton-X for 10 minutes. Cells were then blocked with 5% v/v Donkey serum and 1% w/v BSA in PBS for 1 hour, and then incubated with anti-FLAG primary antibody (Sigma, Cat. #F1804) or anti-H3K9me3 antibody (Abcam Cat. #ab8898) at 4C overnight. Primary antibody was washed off and anti-mouse secondary antibody and Hoescht was added and incubated for 1 hour at room temperature. Mounting media was added and cells were imaged on a Leica SP8 confocal microscope using a 63X oil-immersion lens. Images were analyzed using CellProfiler, where the distribution of FLAG or H3K9me3 staining was assessed within the nucleus (defined by Hoescht staining as a primary object) using MeasureObjectIntensityDistribution with 20 bins. Each cell was considered to be a biological replicate.

### *In vivo* xenotransplantation

All mouse work was carried out in compliance with the ethical regulations approved by the animal research ethics board (AREB) at University Health Network. 8-14 week old immunocompromised mice, NSG (RRID: IMSR_JAX:005557) and NSG-SGM3 (RRID: IMSR_JAX:013062), were sublethally irradiated with 315 cGy 1 day prior to intrafemoral transplantation of HSPCs. NSG-W41 mice were not irradiated prior to transplant. For primary transplants, the input was 10,000 transduced cord blood Lin^−^CD34^+^38^−^HSPCs per mouse. For secondary transplants, the humanCD45^+^mouseCD45^−^BFP^+^ transduced human graft was FACS sorted and >300,000 cells were transplanted per secondary recipient. For secondary transplants at limiting dilution, NSG-SGM3 mice were transplanted with FACS-sorted humanCD45^+^mouseCD45^−^BFP^+^ cells from pooled primary bone marrow at a range of doses (2×10^4^ to 5×10^5^ cells). Positive engraftment was defined as a >0.1% humanCD45^+^mouseCD45^−^ human graft exhibiting both myeloid (CD33^+^) and lymphoid (CD19^+^) populations at 8 weeks post-transplant, in either the transplanted femur or the pooled bone marrow. Frequency of engrafting HSCs was estimated using the *elda* R package.

### RNA-sequencing

10,000-40,000 transduced viable cells were sorted at 3-days post-transduction and RNA was extracted using the PicoPure RNA isolation kit (ThermoFisher, Cat. #KIT0204). RNA-seq libraries were prepared using the NEB Ultra II Directional PolyA kit or NEB Single-cell/Low Input prep kit and sequenced to a depth of 30-60M paired-end reads. Sequences were adapter-trimmed using TrimGalore and aligned to hg38 using STAR on two-pass mode, followed by quantification using Featurecounts (RRID: SCR_012919). Differential expression analysis was performed using DESeq2 (RRID: SCR_000154). Over-representation analyses were performed using g:Profiler and Gene Set Enrichment Analyses were performed using the *fgsea* R package using the Bader Lab geneset collection^71^. Splicing analysis was performed using rMATS^25^. Analysis of previously published RNA-seq data was performed using the same analysis pipeline.

### CUT&RUN-sequencing

MBD1 isoform CUT&RUN was performed as described in Skene et al. (2018) (dx.doi.org/10.17504/protocols.io.zcpf2vn) using 200 000 sorted cells overexpressing FLAG-tagged MBD1 isoforms^72^. The pulldown was performed using mouse anti-FLAG M2 (Sigma, Cat. #F1804, RRID: AB_2910145), with mouse IgG as the isotype control. For histone CUT&RUN, the pulldown was instead performed using rabbit anti-H3K9me3 (Abcam Cat. #ab8898, RRID: AB_306848) and anti-H3K4me3 (C42D8, Cell Signaling Technologies Cat. #9751T), with rabbit IgG as the isotype control. DNA was purified using the Nucleospin PCR cleanup kit (Takara, Cat. # 740609.50) and the library was prepared using NEB Single-cell/Low Input prep kit and sequenced to a depth of 80M paired-end reads. Sequences were adapter-trimmed using TrimGalore and aligned to hg38 using Bowtie2 using the following parameters: –end-to-end –very-sensitive – dovetail –no-mixed –no-discordant –phred33 -I 10 -X 700. Peak-calling was performed using MACS using a threshold of q = 1e-5 to generate consensus peaksets for MBD1-L and MBD1-S. Differential binding analysis was then performed on the combined peakset using DiffBind (RRID: SCR_012918). Metagene density plots and MBD1/histone density were calculated using deepTools.

### ATAC-sequencing

100 000 sorted viable cells were pelleted at 500 x g for 5 minutes at 4C and resuspended in 50 μL of fully supplemented ATAC-seq resuspension buffer (RSB) containing 10mM Tris-HCl (pH 7.4), 10mM NaCl, 3mM MgCl2, 0.1% NP-40, 0.1% Tween-20, and 0.01% digitonin, and water. The cells were incubated on ice for 3 minutes for cell lysis. The lysis was quenched with 1mL of ATAC-seq RSB containing 0.1% Tween-20 (without NP-40 or digitonin). Nuclei were then isolated by centrifugation at 500 x g for 10 minutes at 4C, followed by careful aspiration of supernatant and resuspension of pellet with 50 μL of transposition master mix containing 25ul 2x TD buffer, 2.5 μL transposase (Illumina, cat. no. 200034197), 16.5 μL PBS, 0.5 μL 1% digitonin, 0.5 μL 10% Tween-20, and 5 μL water. Transposition reactions were incubated at 37C for 30 minutes in a thermomixer with shaking at 1000 rpm. Reactions were cleaned up with MinElute columns (Qiagen, cat. No. 28204), followed by library amplification, where PCR cycles were determined by qPCR as described previously^73^. The library was purified and size selected using AMPure XP Beads (Beckman Coulter, catalog ## A63882) and underwent QC by qPCR to confirm enrichment of open chromatin regions (KAT6B and GAPDH) versus closed regions (QML and SLC). Each sample was sequenced to a depth of 120M paired-end reads.

Sequences were adapter-trimmed using TrimGalore and aligned to hg38 using Bowtie2 using the following parameters: -X 2000 –very-sensitive –no-unal. Forward and reverse reads were shifted by 4 and 5 bp respectively to account for Tn5 transposase’s asymmetric cut pattern. Peak-calling was performed using MACS using a threshold of q = 0.01 to generate consensus ATAC-accessible regions for MBD1-L and MBD1-S. Differential accessibility analysis was then performed on the combined peakset using DiffBind.CpG and mCpG density was estimated for each promoter ATAC peak using methylation fractions called from BLUEPRINT methylome CD34+ HSPC bisulfite sequencing data as well as in-house enzymatic methyl-seq data (unpublished)^39^.

### Rapid immunoprecipitation mass spectrometry of endogenous proteins (RIME)

RIME was performed as described in Mohammed et. al. (2016)^40^. Briefly, 15 million transduced MDS-L cells overexpressing MBD1-S or MBD1-L with a C-terminus 3X FLAG tag were cross-linked with 2mM DSG for 20 minutes, then 1% formaldehyde for 10 minutes, and quenched with a 1:10 dilution of 1.25M glycine (pH 7.5) for 5 minutes. Cells were washed twice with ice cold PBS and once with ice cold PBS plus protease inhibitor prior to lysis. Pierce Protein A/G magnetic beads (ThermoFisher ## 88803) were used for immunoprecipitation of the FLAG epitope, where blocked beads were first resuspended in Lysis buffer 3 with 1% Triton X-100 and used to pre-clear the lysates for 1 hour at room temperature. 10 ug of mouse anti-FLAG antibody (Sigma F1804-1MG) was added to the pre-cleared lysate and incubated overnight at 4C. The following day, blocked beads were added to the lysate and incubated at room temperature for 1 hour with mixing. Beads were magnetically separated and washed in 1 mL LiCl RIPA buffer 10 times, and then washed twice with 100mM ammonium hydrogen carbonate (AMBIC) solution. Beads were ensured to be dry, snap frozen and stored at −80C prior to proteomics submission. A volume of 10 μL trypsin solution (20 ng/μl) (Pierce) in 100 mM AMBIC was added to the beads followed by overnight incubation at 37 °C. A second digestion step was performed the next day for 4 h. After proteolysis the tubes were placed on a magnet and the supernatant solution was collected after acidification through step-wise addition of 1uL of 10% TFA to acidify the buffer once it has been separated from the beads.

Then PASEF-DIA LC-MS acquisition was performed. Peptides were loaded onto EvoTip Pure tips for desalting and as a disposable trap column for nanoUPLC using an EvoSep One system. A pre-set EvoSep 100 SPD gradient (from Evosep One HyStar Driver 2.3.57.0) was used with an 8 cm EvoSep C_18_ Performance column (8 cm x 150 μm x 1.5 μm).

The nanoUPLC system was interfaced to a timsTOF HT mass spectrometer (Bruker) with a CaptiveSpray ionisation source (Source). Positive PASEF-DDA, ESI-MS and MS^2^ spectra were acquired using Compass HyStar software (version 6.2, Bruker). Instrument source settings were: capillary voltage, 1,500 V; dry gas, 3 l/min; dry temperature; 180°C. Spectra were acquired between *m/z* 100-1,700. TIMS settings were: 1/K0 0.6-1.60 V.s/cm^2^; Ramp time, 100 ms; Ramp rate 9.42 Hz. Data dependant acquisition was performed with 10 PASEF ramps and a total cycle time of 1.17 s. An intensity threshold of 2,500 and a target intensity of 20,000 were set with active exclusion applied for 0.4 min post precursor selection. Collision energy was interpolated between 20 eV at 0.6 V.s/cm^2^ to 59 eV at 1.6 V.s/cm^2^.

### meRIP peak enrichment at DRACH motifs and E11 splicing reporter quantifications

meRIP enrichment at DRACH motif sites was calculated using the exomepeak R package^55^. Reporter RT-PCR quantification of the long-short ratio was performed using densitometry with ImageJ (RRID: SCR_003070), with a correction factor applied due to differences in band intensities arising from different product lengths.

### Statistical Analysis

All statistical analysis was either performed using GraphPad Prism (GraphPad Software version 10.2.3) or R. A p-value < 0.05 was used as the cutoff for statistical significance. (* p.val. ≤ 0.05, ** p.val. ≤ 0.01, *** p.val. ≤ 0.001).

---

**Supplementary Figure 1:**
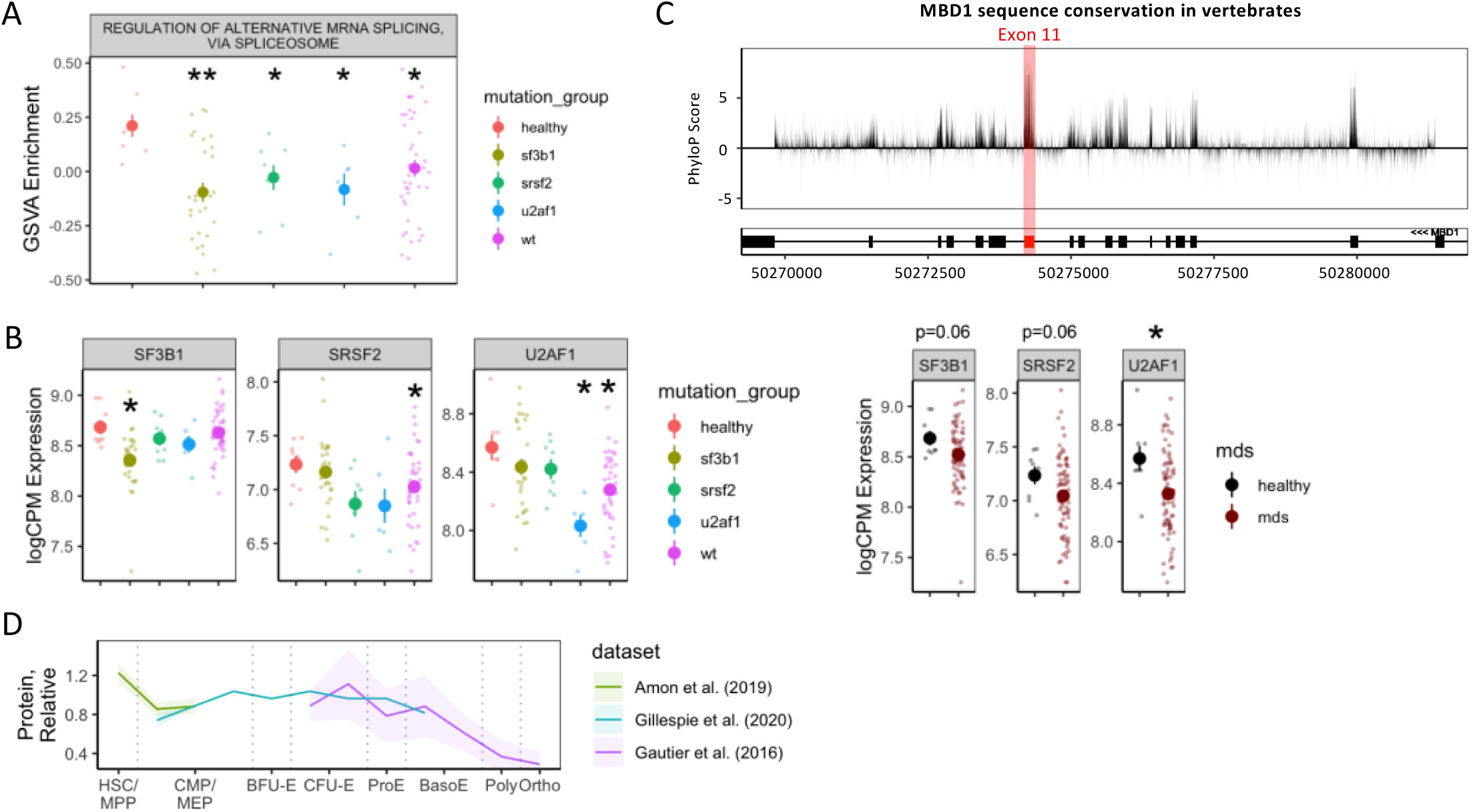
A CXXC-3 containing isoform of MBD1 is globally upregulated in MDS. **A** Pathway enrichment of splicing regulators in splice factor mutant or wildtype MDS subgroups. **B** Expression of SF3B1, SRSF2 and U2AF1 in MDS stratified by mutation status, or between healthy and MDS HSPCs (Pellagatti et. al. 2018 - 8 healthy controls and 82 MDS samples; 28 SF3B1 mutant, 8 SRSF2 mutant, 6 U2AF1 mutant and 40 splice factor wildtype). **C** PhyloP prediction of sequence conservation in MBD1 across vertebrates. Positive scores correspond to conservation whereas negative scores correspond to fast evolution. The magnitude of the score is the −log p-value under the null hypothesis of neutral evolution. **D** Relative expression of MBD1 protein in the hematopoietic hierarchy. (* p.val. ≤ 0.05, ** p.val. ≤ 0.01, *** p.val. ≤ 0.001)

**Supplementary Figure 2:**
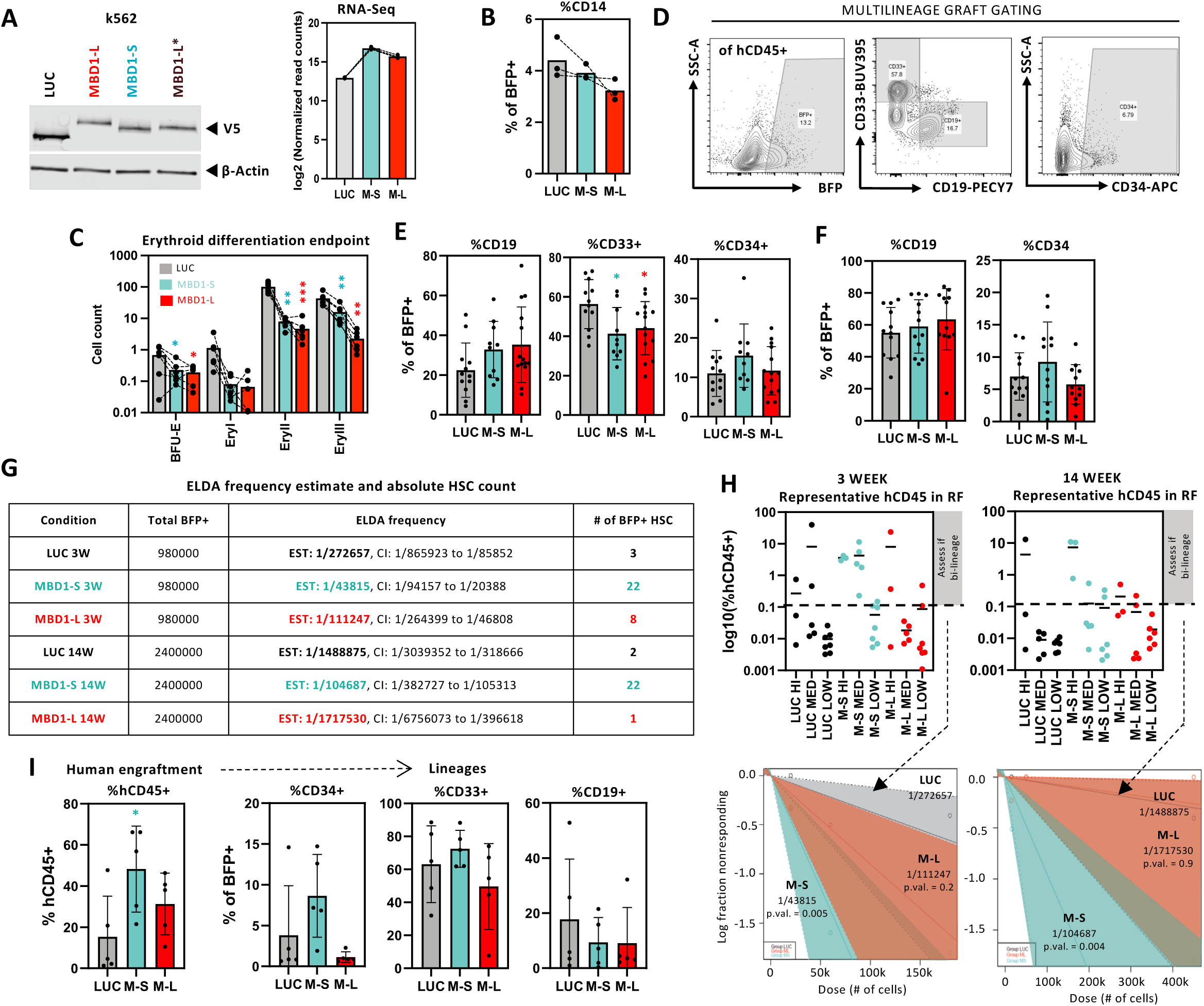
MBD1-L impairs the response to hematopoietic demand in an isoform-specific manner. **A** Western blot validation of lentiviral overexpression of the inert transgene firefly luciferase or various MBD1 isoforms in k562 cells, as well as RNA-based validation of elevated total MBD1 levels in CB CD34+ HSPCs. **B** CD14 expression in cultured Lin-CD34+ HSPCs overexpressing MBD1-L and MBD1-S. **C** Absolute count of erythroid populations at the endpoint of a 2-stage *in vitro* erythroid differentiation culture (N=6 samples). **D** Representative human xenograft gating strategy. **E** Proportion of CD19, CD33 and CD34 within the injected femur of primary 3-week NSG mice transplanted with Lin-CD34+38-HSPCs expressing luciferase or MBD1 isoforms. **F** Proportion of CD19 and CD34 in the injected femur of primary 14-week NGS mice. **G** ELDA calculated frequency, confidence intervals and back-calculated absolute HSC count in pooled primary BFP+ bone marrow from 3- and 14-week primary NSG mice. **H** Human CD45 engraftment at each transplanted dose for 3- and 14-week LDAs. **I** Proportion of human CD45, CD34, CD33 and CD19 in non-limiting secondary transplants of primary 14-week grafts into NSG-SGM3 mice. N=3 samples for *in vitro* assays unless stated otherwise. All transplant data shown is from analysis of the injected femur unless stated otherwise. (* p.val. ≤ 0.05, ** p.val. ≤ 0.01, *** p.val. ≤ 0.001). All statistical tests are performed against Luciferase unless otherwise indicated.

**Supplementary Figure 3:**
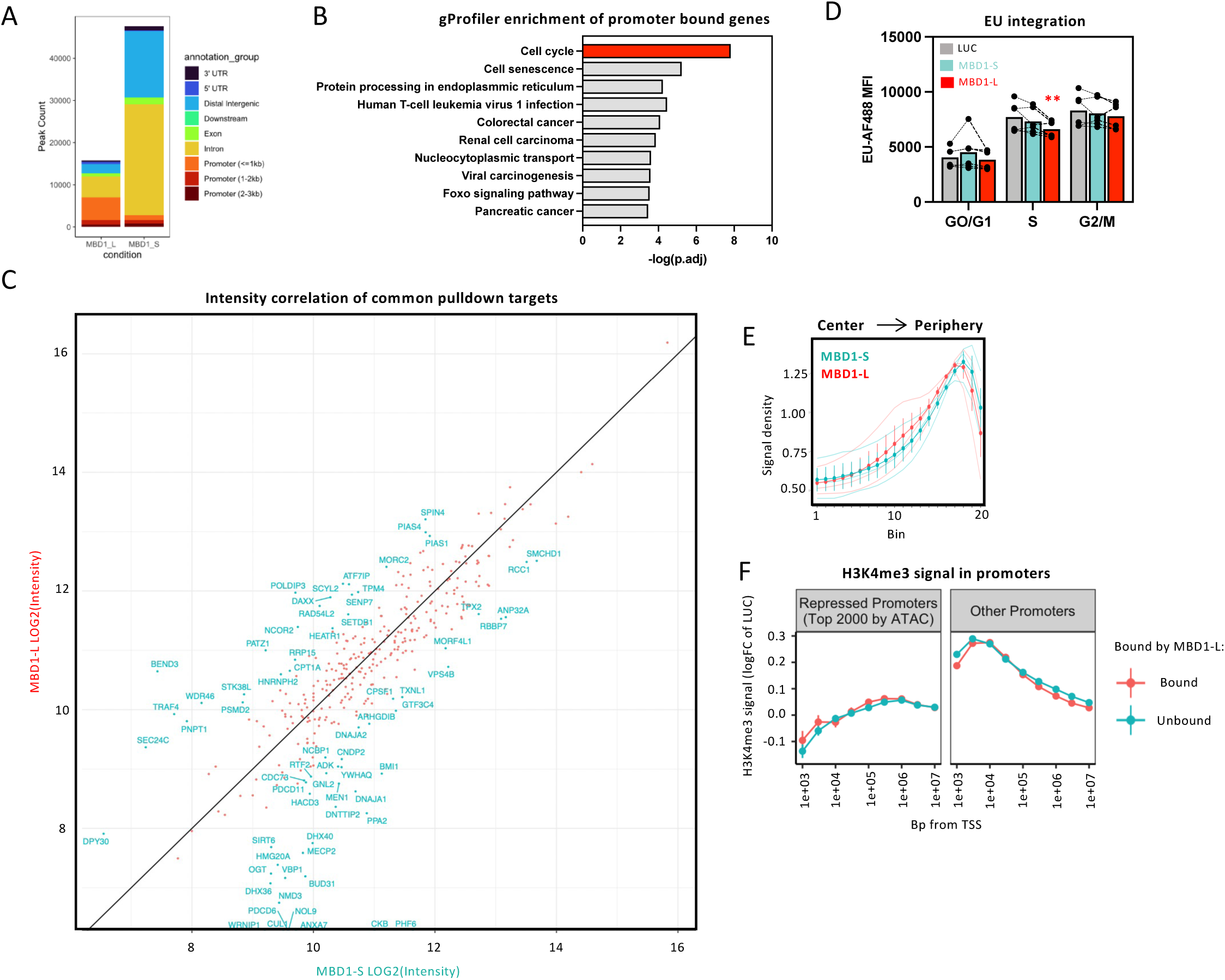
MBD1-L broadly represses promoter CpG islands. **A** MBD1-S and MBD1-L Lin-CD34+ CUT&RUN peak distribution across various functional genomic elements. **B** Top 10 significant gProfiler enrichment terms of MBD1-L promotor bound genes. **C** Correlation of signal intensities of RIME immunoprecipitated MBD1-L and MBD1-S interacting proteins in MDSL cells. **D** Intracellular flow cytometry of MBD1 isoform overexpressing Lin-CD34+ HSPCs pulsed with EU for 1 hour, measured 72 hours post-transduction (N=6 samples). EU signal is plotted against cell cycle phase, annotated via DNA content measured by DAPI signal. **E** Nuclear distribution of H3K9me3 in Lin-CD34+ HSPCs overexpressing MBD1-L or MBD1-S, measured 72 hours post-transduction (N = 30 analyzed cells per sample, with 3 samples used). **F** H3K4me3 CUT&RUN signal proximal to the TSS at MBD1-L bound and repressed promoters compared to luciferase. All multi-omics experiments were performed with 3 independent CB samples. (* p.val. ≤ 0.05, ** p.val. ≤ 0.01, *** p.val. ≤ 0.001)

**Supplementary Figure 4:**
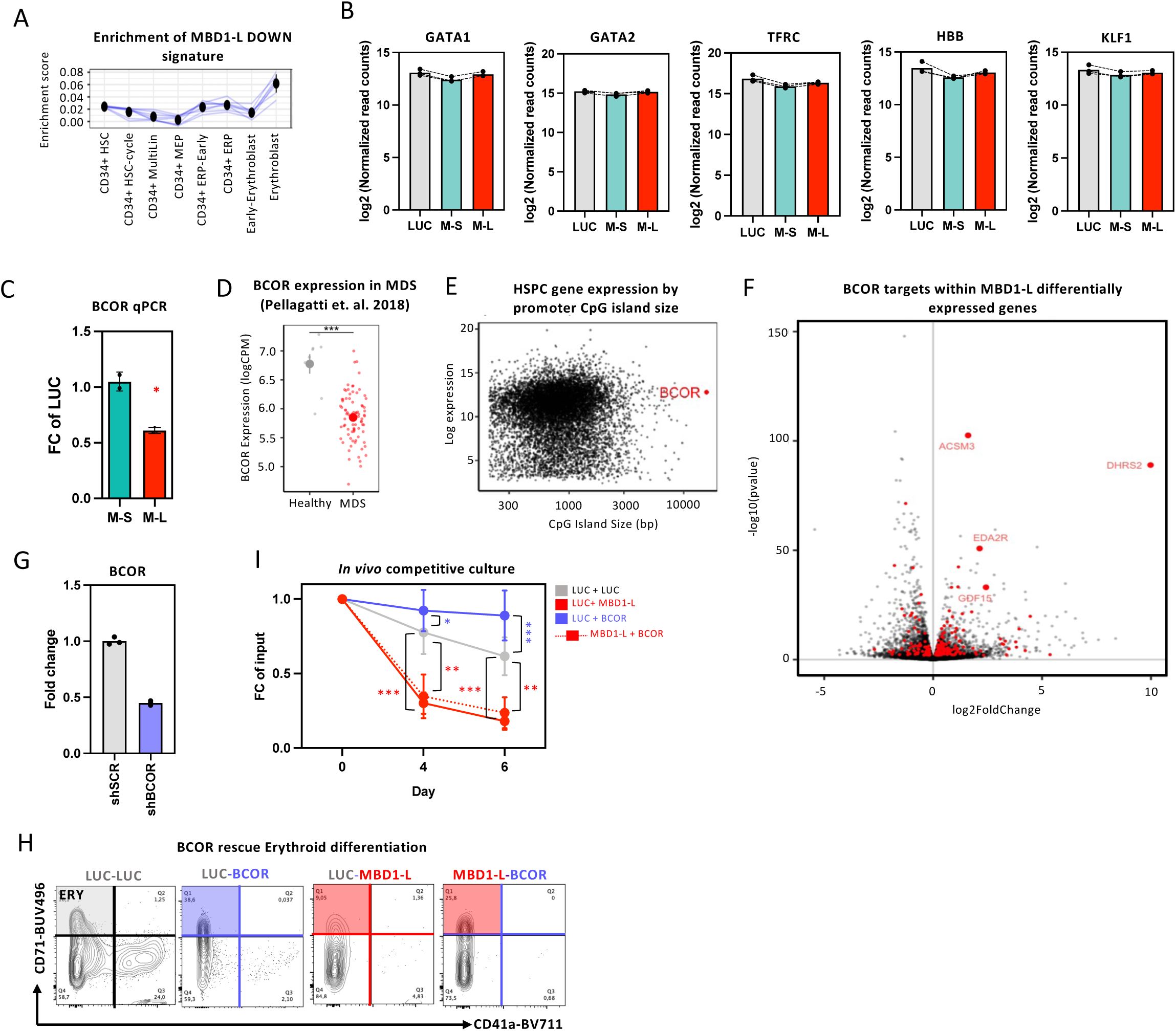
MBD1-L silences promoter networks to inhibit cycling and erythroid differentiation. **A** Mapping of MBD1-L downregulated genes onto stages of erythroid differentiation, data from Hay et al. (2018). **B** Expression of core erythroid genes upon luciferase, MBD1-S or MBD1-L overexpression in Lin-CD34+ HSPCs, plotted from MBD1 isoform RNA-seq. **C** qPCR demonstrating BCOR downregulation after MBD1-L and MBD1-S overexpression. **D** BCOR expression in healthy and MDS CD34+ HSPCs (Pellagatti et al 2018). **E** BCOR CpG island size compared to all other protein coding genes. Compiled through data from UCSC genome browser. **F** Distribution of BCOR-controlled genes within the MBD1-L RNA-seq compared to luciferase. **G** qPCR validation of BCOR targeting shRNA (N=3 technical replicates). **H** Representative flow plots showing rescue of MBD1-L erythroid defects upon co-expression of BCOR. **I** *In vitro* competitiveness of cultured Lin-CD34+ HSPCs transduced with combinations of luciferase, MBD1-L or BCOR (n=6 samples). All multi-omics experiments were performed with 3 independent CB samples. All functional assays are performed with 3 samples unless otherwise stated. (* p.val. ≤ 0.05, ** p.val. ≤ 0.01, *** p.val. ≤ 0.001).

**Supplementary Figure 5:**
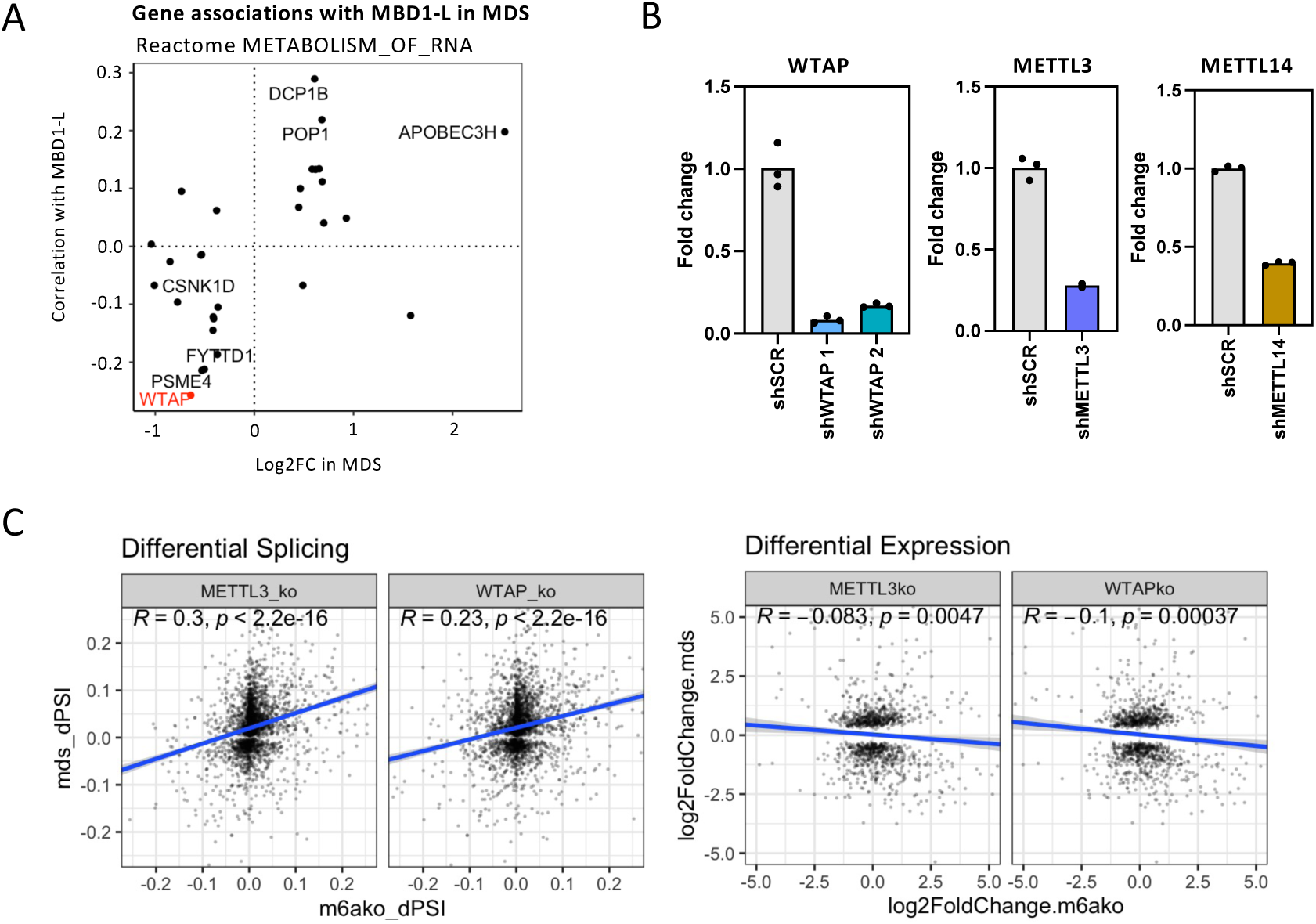
The m6A writer WTAP regulates the splicing of MBD1-L. **A** Correlation between RNA regulators differentially expressed in MDS HSPCs (padj < 0.05) and MBD1-L splicing. The Reactome METABOLISM_OF_RNA geneset was used to filter for splicing regulators. **B** qPCR validation of WTAP, METTL3 and METTL14 shRNAs (N=3 technical replicates). **C** Correlation of splicing signatures between MDS and CRISPR knockout of m6A writers.

**Supplementary Figure 6:**
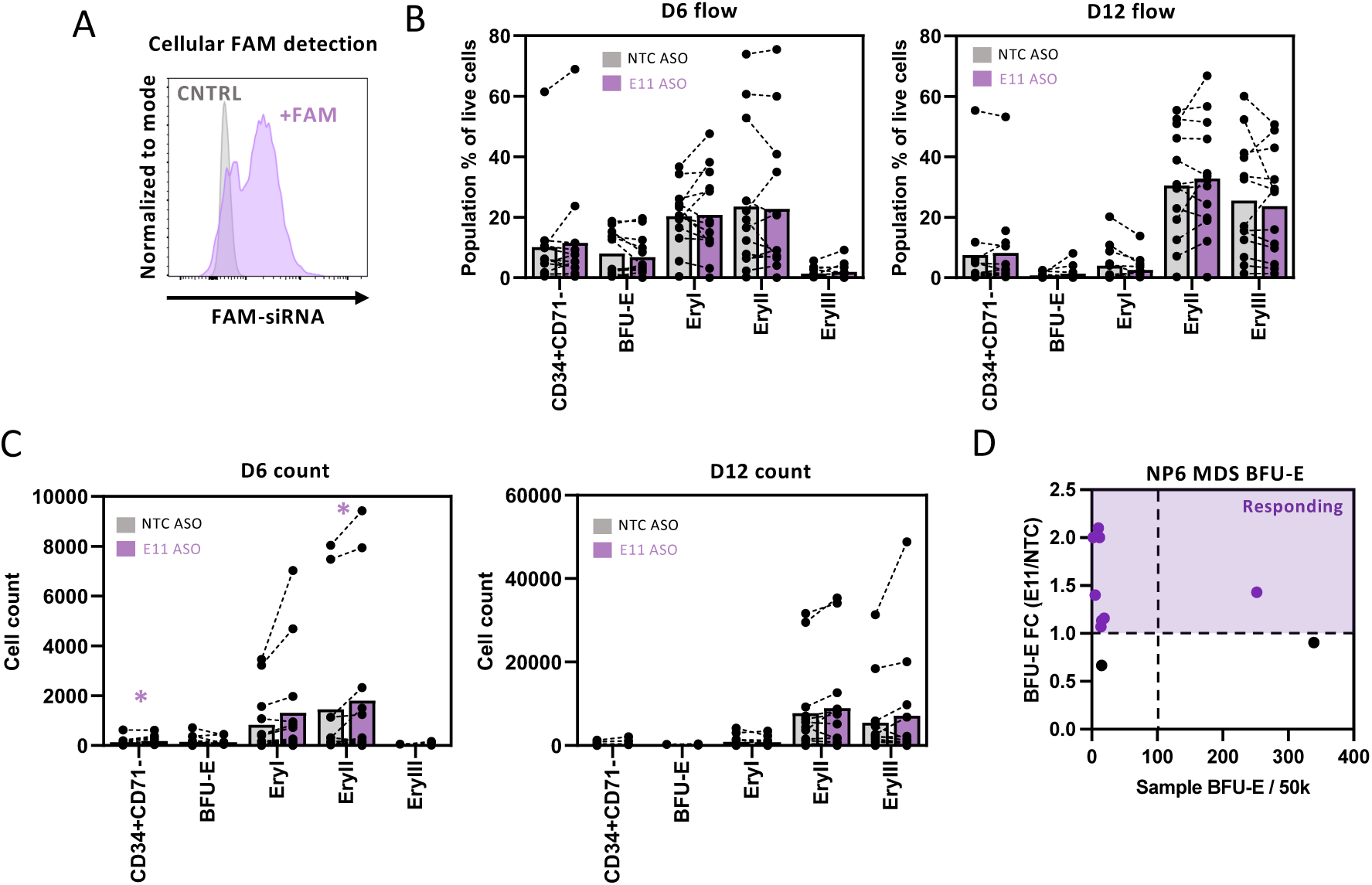
Depletion of MBD1 restores differentiation in MDS cells. **A** CB CD34+ HSPCs treated with either an unlabeled control RNA, or a control RNA conjugated to FAM, where FAM fluorescence is indicative of cellular uptake of the LNPs. **B** Proportion of erythroid populations in primary MDS CD34+ HSPCs treated with E11 or control ASOs at day 6 and 12 of a 2-step erythroid differentiation assay with MDS HSPCs cultured on a primary human bone marrow mesenchymal stromal cell layer (N=13 samples). **C** Absolute count of erythroid populations in primary MDS CD34+ HSPCs treated with E11 or control ASOs at day 6 and 12 of a 2-step erythroid differentiation assay (N=13 samples). **D** BFU-E colonies of primary MDS cells pulsed with E11 and control ASO packaged in a less efficacious LNP formulation with a N/P ratio of 6. Fold change in BFU-E colonies compared to the NTC control are plotted against total BFU-Es in NTC samples. Samples with a BFU-E fold change greater than one are highlighted as responding samples (N=10 samples). (* p.val. ≤ 0.05, ** p.val. ≤ 0.01, *** p.val. ≤ 0.001).

